# A GH81-type β-glucan-binding protein facilitates colonization by mutualistic fungi in barley

**DOI:** 10.1101/2023.04.12.536646

**Authors:** Alan Wanke, Sarah van Boerdonk, Lisa Katharina Mahdi, Stephan Wawra, Miriam Neidert, Balakumaran Chandrasekar, Pia Saake, Isabel M. L. Saur, Paul Derbyshire, Nicholas Holton, Frank L. H. Menke, Mathias Brands, Markus Pauly, Ivan F. Acosta, Cyril Zipfel, Alga Zuccaro

## Abstract

Cell walls are important interfaces of plant-fungal interactions. Host cell walls act as robust physical and chemical barriers against fungal invaders, making them an essential line of defense. Upon fungal colonization, plants deposit phenolics and callose at the sites of fungal penetration to reinforce their walls and prevent further fungal progression. Alterations in the composition of plant cell walls significantly impact host susceptibility. Furthermore, plants and fungi secrete glycan hydrolases acting on each other’s cell walls. These enzymes release a wide range of sugar oligomers into the apoplast, some of which trigger the activation of host immunity *via* host surface receptors. Recent characterization of cell walls from plant-colonizing fungi have emphasized the abundance of β-glucans in different cell wall layers, which makes them suitable targets for recognition. To characterize host components involved in immunity against fungi, we performed a protein pull-down with the biotinylated β-glucan laminarin. Thereby, we identified a glycoside hydrolase family 81-type glucan-binding protein (GBP) as the major β-glucan interactor. Mutation of *GBP1* and its only paralogue *GBP2* in barley led to decreased colonization by the beneficial root endophytes *Serendipita indica* and *S. vermifera*, as well as the arbuscular mycorrhizal fungus *Rhizophagus irregularis.* The reduction of symbiotic colonization was accompanied by enhanced responses at the host cell wall. Moreover, *GBP* mutation in barley also increased resistance to fungal infections in roots and leaves by the hemibiotrophic pathogen *Bipolaris sorokiniana* and the obligate biotrophic pathogen *Blumeria graminis* f. sp. *hordei,* respectively. These results indicate that GBP1 is involved in the establishment of symbiotic associations with beneficial fungi, a role that has potentially been appropriated by barley-adapted pathogens.

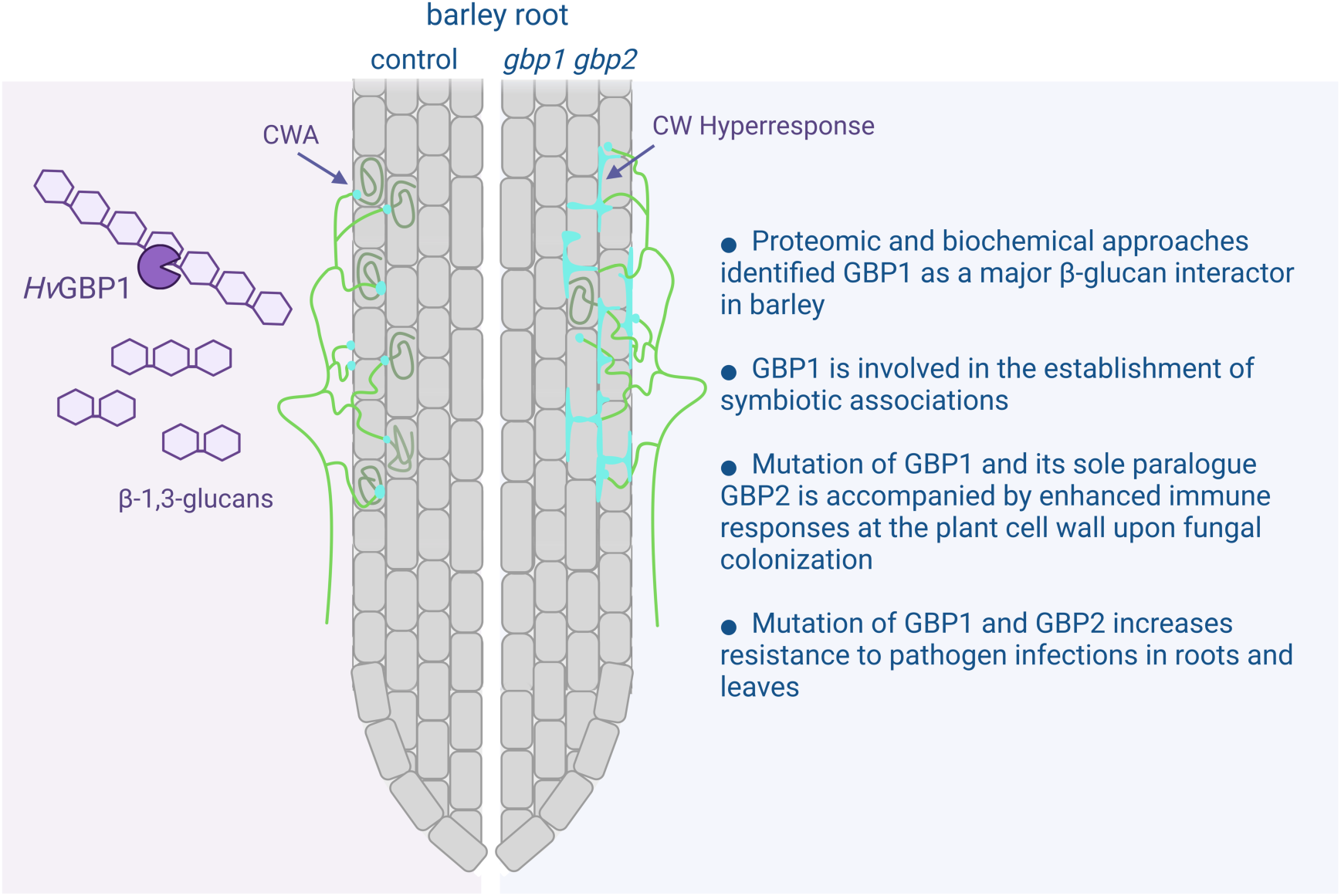

**In Brief:** GBP1, a family 81 glycoside hydrolase, is an important β-glucan interactor in barley. Mutation of GBP1 and its sole paralogue GBP2 leads to reduced colonization by beneficial root endophytes, AM fungi and pathogens, accompanied by enhanced responses at the plant cell wall. This indicates that GBP1 and β-glucans are compatibility factors involved in the establishment of symbiotic associations with beneficial fungi, a role possibly hijacked by pathogens.

## Introduction

Cell walls (CWs) are key interfaces in plant-fungal interactions, representing the first point of physical contact between the invading fungus and its potential host. Plant CWs consist of adaptable networks of cellulose microfibrils embedded in a matrix of hemicellulose (mostly xyloglucan, mannan and xylans), pectins and hydrophobic polymers such as cutin, lignin and suberin. Their architecture and composition are highly responsive to external and internal cues (Cosgrove, 2005; Cosgrove & Jarvis, 2012). Plant CWs act as scaffolds for hydrolytic enzymes and reservoirs for antimicrobial substances such as secondary metabolites and reactive oxygen species (ROS) (Aist, 1977; Brown *et al*., 1998; Bednarek *et al*., 2009; Underwood, 2012). To gain access to the nutritional resources of the host, fungi need to trespass these chemical and physical barriers. They do so by employing either carbohydrate active enzymes (CAZymes) and/or specialized pressure-driven penetration structures like appressoria. Plants counteract those strategies by forming carbohydrate-enriched CW appositions (CWAs), so-called papillae, which act as CW reinforcements at the sites of fungal penetrations (Hückelhoven, 2007). Although the relevance of papillae in preventing or delaying fungal colonization has been controversial in recent decades, many studies correlated the effectiveness of papillae to the timing of deposition, size, architecture and composition of the papillae (Aist, 1977; Zeyen *et al*., 2002; Chowdhury *et al*., 2014; Hückelhoven, 2014). Systematic screenings of *Arabidopsis thaliana* CW mutants have shown that interference with the structure and composition of plant CWs severely impacts fungal compatibility (Molina *et al*., 2021). Besides the direct effects of changes in CW architecture, compromised CW integrity can indirectly prime host defenses and/or alter phytohormone levels, ultimately impacting the success of fungal colonization (Vogel *et al*., 2002; Nishimura *et al*., 2003; Hernández-Blanco *et al*., 2007; Escudero *et al*., 2017; Gavrin *et al*., 2020; Molina *et al*., 2021).

Both fungi and plants secrete a plethora of CAZymes upon confrontation. These host and fungal hydrolases not only physically damage each other’s CWs, but also release carbohydrate oligomers as degradation products that can inform the host’s immune system about the invading microbe. Fungal chitin and its deacetylated derivative chitosan are well studied microbe-associated molecular patterns (MAMPs) that upon perception by cell surface receptors mount a range of defense responses such as the production of ROS and the secretion of antimicrobials (Yin *et al*., 2016; Albert *et al*., 2020). Furthermore, fungal hydrolases and mechanical forces exerted during fungal penetration release a variety of plant CW-derived fragments, functioning as damage-associated molecular patterns (DAMPs) (Hahn *et al*., 1981). This recognition of modified self similarly initiates immune signaling pathways. Examples for unequivocal plant CW-derived DAMPs are cellulose oligomers, pectin oligogalacturonides, as well as mannan, xyloglucan and arabinoxylan fragments (Aziz *et al*., 2007; Galletti *et al*., 2008; Ferrari *et al*., 2013; Claverie *et al*., 2018; Zang *et al*., 2019; Mélida *et al*., 2020, Zarattini *et al*., 2021; Liu *et al*., 2023).

In the context of fungal colonization, β-glucans perform different functions in enhancing or suppressing immune responses, depending on their structure and branch decorations and cannot be clearly categorized as MAMPs or DAMPs. They represent the major component of most fungal cell surface glycans, being part of both the outer layer of the rigid fungal CW as well as the surrounding gel-like extracellular polysaccharide (EPS) matrix loosely attached to the CW (Wawra *et al*., 2019; Wanke *et al*., 2021). Plants secrete various carbohydrate hydrolases, amongst them β-glucanases, that target these fungal glycans (Rovenich *et al*., 2016; Perrot *et al*., 2022). The potential of some glycans to act as MAMPs and elicit pattern-triggered immunity has been demonstrated for a wide range of host plants (Fesel & Zuccaro, 2016). Notably, plants respond differentially to short-chain and long-chain β-1,3-glucans (Wanke *et al*., 2020). While *Arabidopsis thaliana* Col-0 mounts immune responses such as ROS production, cytosolic calcium influx, MAPK activation and PR gene expression induction upon treatment with short-chain β-1,3-glucans like laminarihexaose, it does not respond to long-chain β-1,3-glucans. In contrast, immune responses in *Nicotiana benthamiana* are only activated upon treatment with long-chain β-1,3-glucans. The monocots *Hordeum vulgare* (barley) and *Brachypodium distachyon* are able to perceive both types of β-1,3-glucan independent of their degree of polymerization (Mélida *et al*., 2018; Wanke *et al*., 2020). While the responses to laminarihexaose in Arabidopsis are mediated by the central carbohydrate lysin motif (LysM) receptor-like kinase CERK1, perception of long-chain β-1,3-glucans in *N. benthamiana* and *Oryza sativa* is CERK1 independent (Mélida *et al*., 2018; Wanke *et al*., 2020). It was recently suggested that β-glucan oligomers from fungal cell surface glycans do not function only as MAMPs. Upon fungal colonization, barley secretes the β-1,3-glucanase BGLUII (apoplastic glycoside hydrolase 17 family, GH17) which releases a highly substituted and specific β-1,3-glucan decasaccharide (β-GD) from the fungal EPS matrix (Chandrasekar *et al*., 2022). Instead of activating immune responses in its host, β-GD exhibits antioxidative properties that help to overcome the hostile oxidative environment and, thereby, facilitate host colonization. In addition to their occurrence in fungal CWs, β-1,3-glucan polymers are major components of callose in host CWAs (Hückelhoven, 2007). Since the release of callose fragments from papillae by mechanical force or enzymatic digest is conceivable, β-1,3-glucan oligomers can serve a dual function as DAMPs and/or MAMPs (Mélida *et al*., 2018, Martín-Dacal *et al*., 2023). A similar dual role has been recently attributed to immunogenic mixed-linked β-1,3/1,4-glucans that can be found in the CWs of grasses as well as fungi and other microbes (Rebaque *et al*., 2021; Barghahn *et al*., 2021).

To detect novel components linked to β-glucan signaling in barley, we performed a protein pull-down with biotinylated laminarin, which has the β-1,6-branched β-1,3 glucans also found in fungal cell walls. Thereby, we identified the GH81 β-1,3- endoglucanase GBP1 as a major interactor of β-1,3/1,6-glucans. Purified GBP1 specifically hydrolyzes β-1,3-linked glucans and can modulate ROS responses in *N. benthamiana* and *H. vulgare*. CRISPR/Cas9-generated mutation of the two *GBP* gene copies decreased the colonization efficiency of the beneficial root endophytes *Serendipita indica* and *S. vermifera* (Basidiomycota) as well as the arbuscular mycorrhizal (AM) fungus *Rhizophagus irregularis* (Glomeromycotina). This phenotype was accompanied by a hyperresponse of the host CW during fungal colonization. Additionally, *gbp* double mutants were more resistant to root infection with the hemibiotrophic pathogen *Bipolaris sorokiniana* and leaf infection with *Blumeria graminis* f. sp. *hordei (*hereafter*, B. graminis)* (Ascomycota). Altogether, this demonstrates a previously undescribed role of β-1,3-endoglucanases as tissue-independent broad range compatibility factors for fungal colonization.

## Results

### Barley GBP-type enzymes mediate compatibility to leaf and root colonization by fungi with different lifestyles

Leaf and root tissues of barley express the cellular machinery necessary to sense the long-chain β-1,3/β-1,6 branched glucan laminarin (Supplementary Figure 1). To discover interactors of long-chain β-glucans, we performed a protein pull-down approach using biotinylated laminarin as bait. The biotinylated and non-biotinylated versions of laminarin induced similar ROS burst responses in barley (Supplementary Figure 2A), confirming that biotinylation of laminarin does not alter its immunogenic potential. Non-biotinylated laminarin and biotinylated elf18 were included as control treatments. Elf18 is a peptide derived from the prokaryotic elongation factor Tu, a well characterized immunogenic peptide solely perceived by *Brassicaceae* members (Kunze *et al*., 2004; Zipfel *et al*., 2006). Eluted proteins were separated by SDS-PAGE (Supplementary Figure 2B), extracted from the gel, and further subjected to tandem mass spectrometric analysis. A total of 101 proteins were identified across the three treatments, based on strict criteria for candidate selection (i.e. proteins detected in three out of four replicates). The only interactor detected exclusively in the biotinylated laminarin treatment (Figure 1A) was the β-glucan-binding protein GBP1 (HORVU5Hr1G059430.56). Despite the absence of a predicted signal peptide in its gene sequence, GBP1 was previously identified in barley apoplastic fluids following colonization by the mutualistic root endophyte *S. indica* (Nizam *et al*., 2019). In support of this, another study demonstrates that a GBP ortholog in soybean is secreted into the apoplastic space even in the absence of a signal peptide (Umemoto *et al*., 1997). GBPs are found in the genomes of most land plants, including bryophytes, ferns and angiosperms (Supplementary Figure 3). They have been duplicated several times throughout the evolution of plants, with a large expansion in legumes. Barley has two GBP copies (GBP1 and GBP2, HORVU6Hr1G034610.3) but only GBP1 is expressed according to publicly available transcriptomic datasets and our own expression analyses (Supplementary Figure 4).

**Figure 1.**
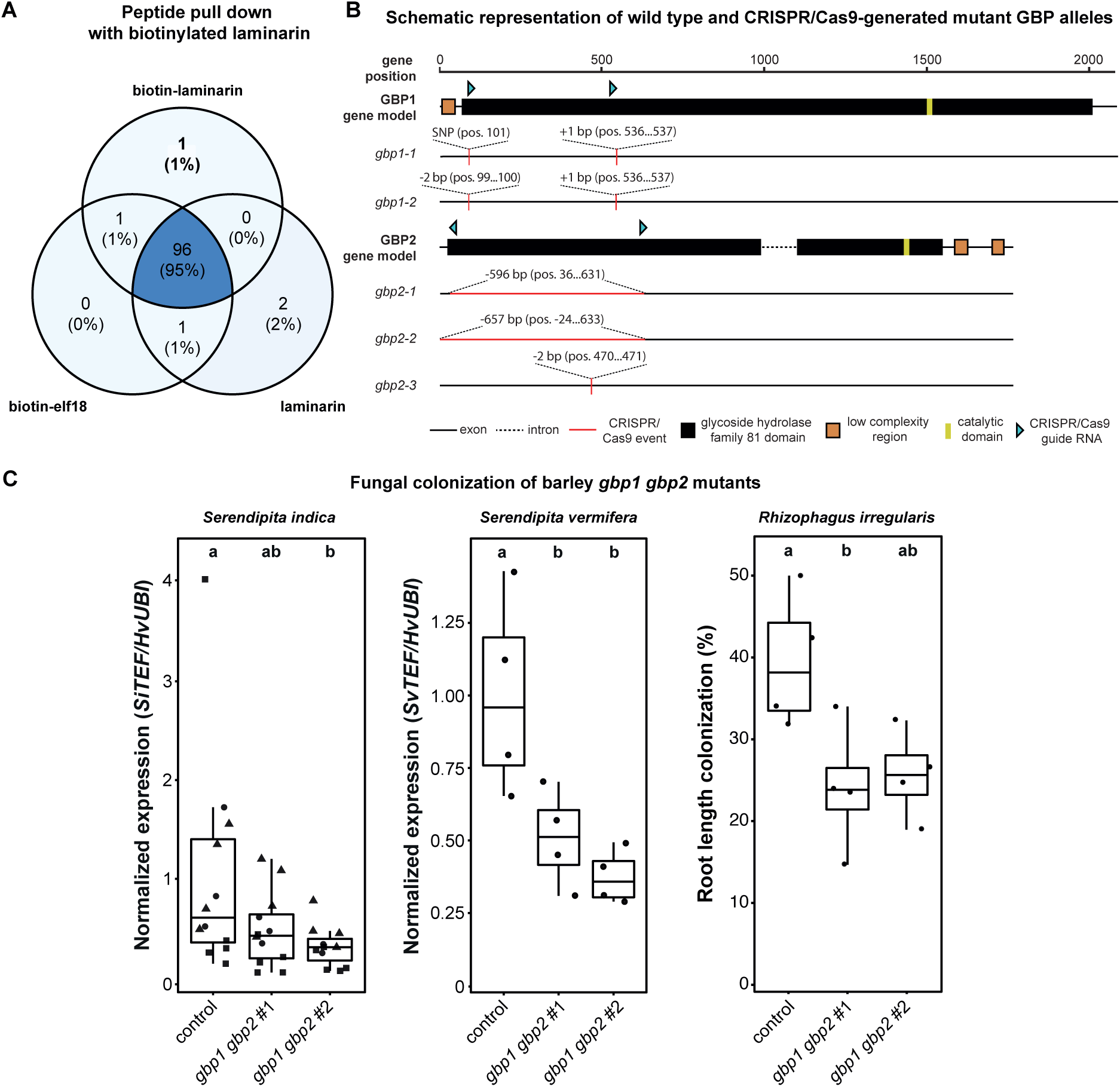
GBP1 is a compatibility factor identified *via* protein pull-down with biotinylated β-glucan in barley. (A) Venn diagram of barley proteins identified by LC-MS/MS analysis with biotinylated β-glucan laminarin, biotinylated elf18 (peptide derived from bacterial elongation factor Tu) or non-biotinylated β-glucan laminarin as bait. Only the proteins whose peptides were identified in three out of four replicates are listed. (B) Schematic overview of CRISPR/Cas9-generated mutant alleles of *GBP1* and *GBP2*. (C) Fungal colonization assays in roots of control and *gbp1 gbp2* mutant lines. The expression of *S. indica* and *S. vermifera* housekeeping genes were quantified by RT-PCR and normalized to the barley housekeeping gene ubiquitin (*Hv*UBI). *R. irregularis* colonization was assessed using light microscopy to quantify the presence of ink-stained *R. irregularis* structures in roots. Root sections were considered to be colonized when arbuscules, intraradical hyphae (IRH) or vesicles were present. A total of 300 randomly chosen sections (covering 30 cm of root) were analyzed per replicate (n=4). Boxplot elements in this figure: center line, median; box limits, upper and lower quartiles; whiskers, 1.5 × interquartile range. Data points from independent experiments are indicated by different shapes. Different letters represent statistically significant differences based on one-way ANOVA and Tukey’s post hoc test (significance threshold: *P* ≤0.05).

To investigate the contribution of barley GBPs to fungal colonization, we used a CRISPR/Cas9 approach to generate plants mutated in both *GBP* genes (*gbp1 gbp2*, Figure 1B). The first mutant line (*gbp1 gbp2* #1) is homozygous for the mutant alleles *gbp1-1* and *gbp2-1*, and the second mutant line (*gbp1 gbp2* #2) is homozygous for the *gbp1-2 allele* and biallelic for *gbp2-2* and *gbp2-3*. The seedlings of the two independent mutant lines showed no differences in root and shoot biomass compared to the control lines (Supplementary Figure 5). To survey colonization by fungi with different lifestyles and colonization strategies, we inoculated the control and both mutant lines with the beneficial root endophytes *S. indica* and *S. vermifera* as well as the AM fungus *R. irregularis* (Figure 1C, Supplementary Figure 6). In all three cases, total fungal colonization was reduced in both *gbp1 gbp2* mutant lines compared to the control lines, with at least one mutant line presenting a significant reduction in colonization. To expand the range of surveyed fungal lifestyles, we additionally tested colonization by the hemibiotrophic root pathogen *Bipolaris sorokiniana* and the obligate biotrophic foliar pathogen *Blumeria graminis* (Supplementary Figure 7). Both *B. sorokiniana* and *B. graminis* showed a significant decrease of fungal biomass or penetration success, respectively. In conclusion, the *gbp1 gbp2* mutant lines showed increased resistance to fungal colonization compared to the control plants, irrespective of the taxonomic position of the fungus, its lifestyle or the host tissue inoculated. Although we cannot rule out the possibility that GBP2 is also involved in mediating compatibility, the fact that we did not detect any expression of GBP2 (Supplementary Figure 4) suggests that the observed phenotype mainly depends on the role of GBP1 as a compatibility factor for fungal colonization of root and leaf tissues.

### GBP1 is an active β-1,3-endoglucanase hydrolyzing β-1,3-glucans with a low degree of β-1,6 substitutions

We further biochemically characterized the β-glucan-interactor GBP1, which belongs to the GH81 family of the carbohydrate active enzymes (CAZymes). Enzymes of this GH-family are characterized by their endoglycosidic hydrolase activity on substituted and unsubstituted β-1,3-linked glucans. Their catalytic center is highly conserved across bacteria, fungi, and plants. GH81 family glycosyl hydrolases follow an inverting hydrolysis mechanism that requires a glutamate residue acting as nucleophile (Fig. 2A).

**Figure 2.**
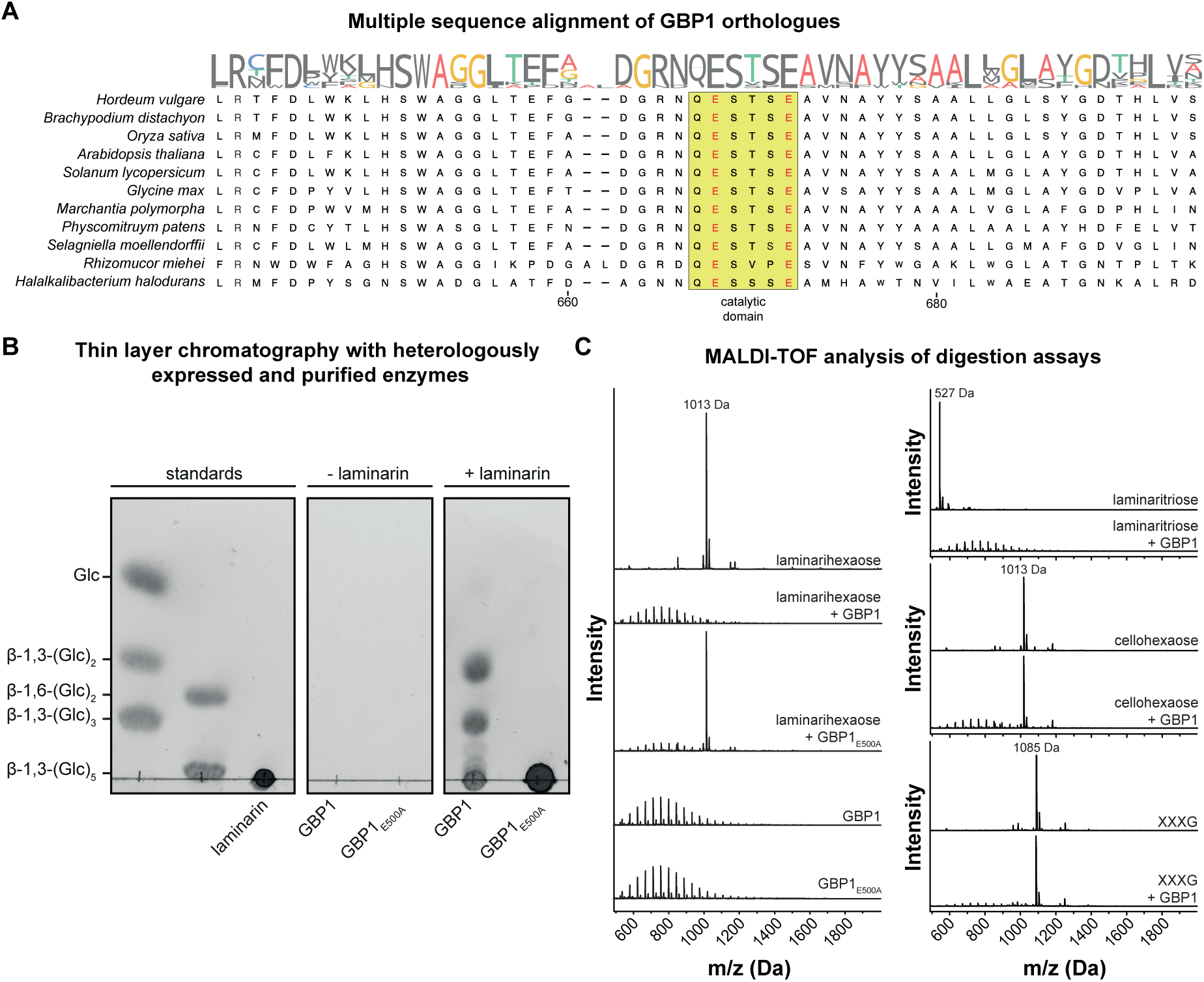
Barley GBP1 has a conserved GH81 catalytic center that specifically mediates the hydrolysis of β-1,3-glucans. (A) Multiple sequence alignment with sequences from different domains of life indicate conservation of the catalytic domain (yellow box), which contains the previously described glutamate residues (red letters) that act as nucleophiles in carbohydrate hydrolysis (Fliegmann *et al*., 2004). (B) Analysis of the β-1,3-glycosidic activity of heterologously expressed and purified GBP1 and mutant GBP1_E500A_ on laminarin from *Laminaria digitata*. After overnight incubation at 25°C, the digestion products were analyzed by thin layer chromatography. The wild-type, but not the mutated version of GBP1, exhibited hydrolytic activity on laminarin. The experiment was repeated at least four times with similar results. (C) The activity of GBP1 and GBP1_E500A_ was tested on laminarihexaose (β-1,3-glucan hexamer), laminaritriose (β-1,3-glucan trimer), cellohexaose (β-1,4-glucan hexamer) and XXXG (xyloglucan heptasaccharide). After overnight digestion at 25°C, the products were analyzed by MALDI-TOF mass spectrometry. Among the tested substrates, the hydrolytic activity of laminarin was found to be specific to β-1,3-glucans. The ion signal ladder between 500-1000 Da represents an unknown contamination present in the GBP1 preparation and is thus unrelated to any digested carbohydrate. Digestion assays were performed twice with similar results. UT, untreated; XXXG, xyloglucan heptasaccharide.

To verify the predicted β-1,3-endoglycosidic activity of GBP1, we overexpressed and purified a codon-optimized, C-terminally HA-StrepII-tagged version of barley GBP1 from transiently transformed *N. benthamiana* leaves (Supplementary Figure 8). Digestion assays with laminarin as a substrate confirmed the β-1,3-endoglycosidase activity of GBP1 (Figure 2B). Furthermore, site-directed mutagenesis of the predicted catalytically active site of GBP1 by replacing the first glutamate residue (putative nucleophile, E500) with an alanine (GBP1E500A) abolished its enzymatic activity (Figure 2B). This confirms that the observed digestion of laminarin is specific to the conserved catalytic site of GBP1. Activity assays with laminarin showed that GBP1 is highly active over a wide range of pH values (pH 5-9) and has the highest hydrolytic activity at 60°C (Supplementary Figure 9). GBP1 activity is specific to β-1,3-linked glucans (laminarihexaose, laminaritriose and laminarin), while MALDI-TOF analysis revealed that GBP1 is unable to digest other oligosaccharides such as β-1,4-linked cellohexaose and xyloglucan (XXXG) (Figure 2B and 2C).

To further test whether GBP1 is able to process complex substrates derived from the CW and extracellular polysaccharide (EPS) matrix of fungi, we performed digestion assays with crude *S. indica* CW and EPS matrix preparations as well as the previously characterized β-GD (Chandrasekar *et al*., 2022). β-GD is a glucan fragment that is highly enriched during digestion of the *S. indica* matrix with the barley glucanase BGLUII. While GBP1 does not hydrolyze the highly β-1,6-branched glucans in the EPS matrix of *S. indica* and the derived β-GD, it releases oligosaccharides (DP 7-13) from *S. indica* CWs to a limited extent (Supplementary Figure 10). This is consistent with the inability of GBP1 to digest the highly substituted laminarin from the brown alga *Eisenia bicyclis* (Supplementary Figure 11). This verifies that frequent β-1,6-branching patterns of glucans protect against hydrolytic degradation and activation of plant immunity as previously demonstrated for the EPS matrices of plant-colonizing fungi (Chandrasekar *et al*., 2022).

Secreted plant hydrolases are known to modulate the activation of immunity by releasing, tailoring and increasing the accessibility of fragments from invading microbes (Rovenich *et al*., 2016; Buscaill *et al*., 2019). To test to what extent the observed hydrolytic activity of GBP1 on β-1,3-glucans alters their immunogenic potential, we treated *N. benthamiana* and barley with gradually digested laminarin (Figure 3A and 3B).

**Figure 3.**
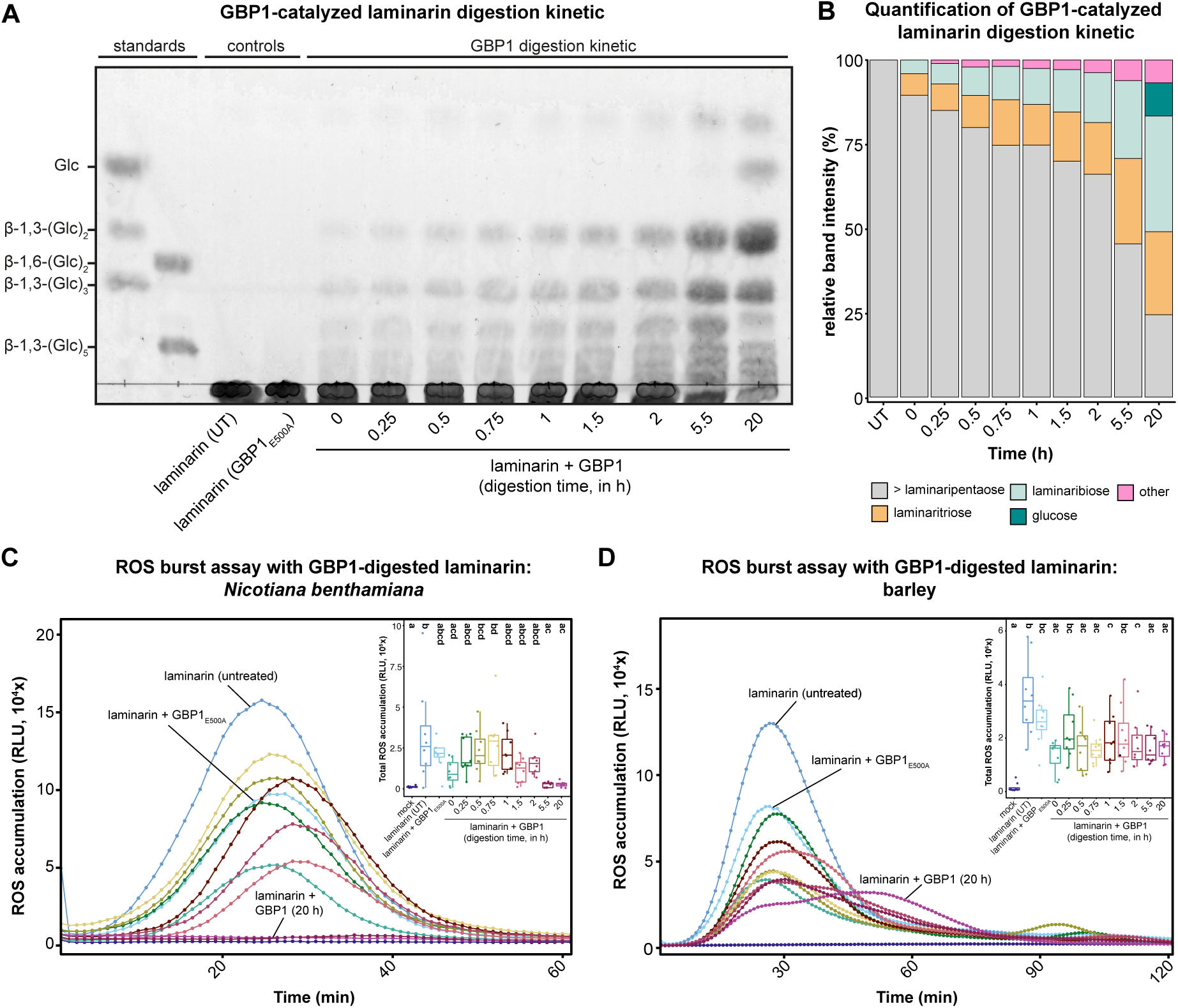
GBP1 treatment of long-chain β-1,3-linked glucans modulate ROS production. To investigate the effect of gradual hydrolysis of β-glucans by GBP1 on ROS accumulation in *N. benthamiana* and barley, a digestion time series was performed with laminarin. (A) The digestion products were analyzed by thin layer chromatography (TLC). Untreated (UT) and GBP_E500A_-treated laminarin (20 hours) served as controls. Digestion was performed at 25°C. (B) Quantification of GBP1-catalyzed laminarin digestion products. TLC band intensity was quantified using the ImageJ software. (C and D) Apoplastic ROS accumulation after treatment of *N. benthamiana* (C) and barley (D) with gradually digested laminarin (1 mg·ml^-1^) was monitored using a luminol-based chemiluminescence assay. Milli-Q water (mock) treatment, untreated laminarin (UT) and GBP1_E500A_ were used as controls. Values represent mean ± SEM from eight replicates. Boxplots display total ROS accumulation over the measured period of time. The assays were performed two times with independent laminarin digestions. Boxplot elements in this figure: center line, median; box limits, upper and lower quartiles; whiskers, 1.5 × interquartile range. Different letters represent statistically significant differences in expression based on a one-way ANOVA and Tukey’s post hoc test (significance threshold: *P* ≤0.05). ROS, reactive oxygen species. RLU, relative luminescence units.

To assess changes in early immune signaling, we used β-glucan-triggered ROS production, one of the first immunity outputs triggered by MAMPs (Yu *et al*., 2017), as a readout. *N. benthamiana* and barley differ in terms of their perceived β-glucan substrates (Wanke *et al*., 2020). The dicotyledonous *N. benthamian*a activates immune responses only when treated with long-chain β-1,3-glucans. As digestion of laminarin by GBP1 progressed, the intensity of the triggered oxidative burst gradually decreased to the level of mock treatment (20 hours) (Figure 3C). In contrast to *N. benthamiana*, barley responds to both short- and long-chain β-1,3-glucans. Accordingly, incubation of laminarin with GBP1 reduced the intensity of the oxidative burst but did not completely abolish it due to the recognition of the released short-chain hydrolysis products (Figure 3D). Treatment with the inactive version GBP1E500A (20 hours) did not affect laminarin perception in either species (Figure 3C and 3D).

### Barley GBPs are not essential for β-glucan-triggered ROS production

Previous studies on perception of a β-1,3/1,6-heptaglucoside from oomycetes in legumes have led to the suggestion that GBPs may be part of the corresponding β-glucan perception machinery (Fliegmann *et al*., 2004; Umemoto *et al*., 1997). Since oomycete-derived heptaglucoside perception is specific to legumes (Fliegmann *et al*., 2004), we investigated whether barley GBPs are involved in β-glucan-triggered ROS production upon treatment with short-and long-chain β-glucans. Since ROS production still occurred in both *gbp* double mutant lines (Supplementary Figure 12), we conclude that GBPs are not essential for the immune perception of β-glucan in barley.

### Barley GBP mutants exhibit hyperresponsive CW reaction upon colonization by endophytic fungi

Previous studies have shown that broad-spectrum resistance to fungi can be mediated by the enhanced formation of CWAs, so called papillae, which are predominantly formed at the sites of hyphal penetration (Hilbert et al., 2019). A major component of these CW reinforcements is callose, a linear β-1,3-linked glucan that could serve as a substrate for GBPs. To investigate the effect of GBPs on the formation of CWAs upon colonization by the root endophytes *S. indica* and *S. vermifera*, we monitored and quantified CWAs using a fluorescent version of the carbohydrate-binding lectin concanavalin A (ConA), which has been shown to accumulate at the CWAs in barley in previous studies (Zuccaro *et al*., 2011; Hilbert *et al*., 2019).

Papilla formation was observed in all barley lines tested after colonization with the fungus, with no differences in papilla size or frequency between the control line and the two *gbp1 gbp2* mutant lines (Supplementary Figure 13). However, in addition to round-shaped papillae, inoculated mutant roots exhibited an extensive CW response lining the CWs of the epidermal root cells (Figure 4). This response occurred exclusively when colonized by the fungus and not in plants grown without the fungus under control conditions (Supplementary Figure 14).

**Figure 4.**
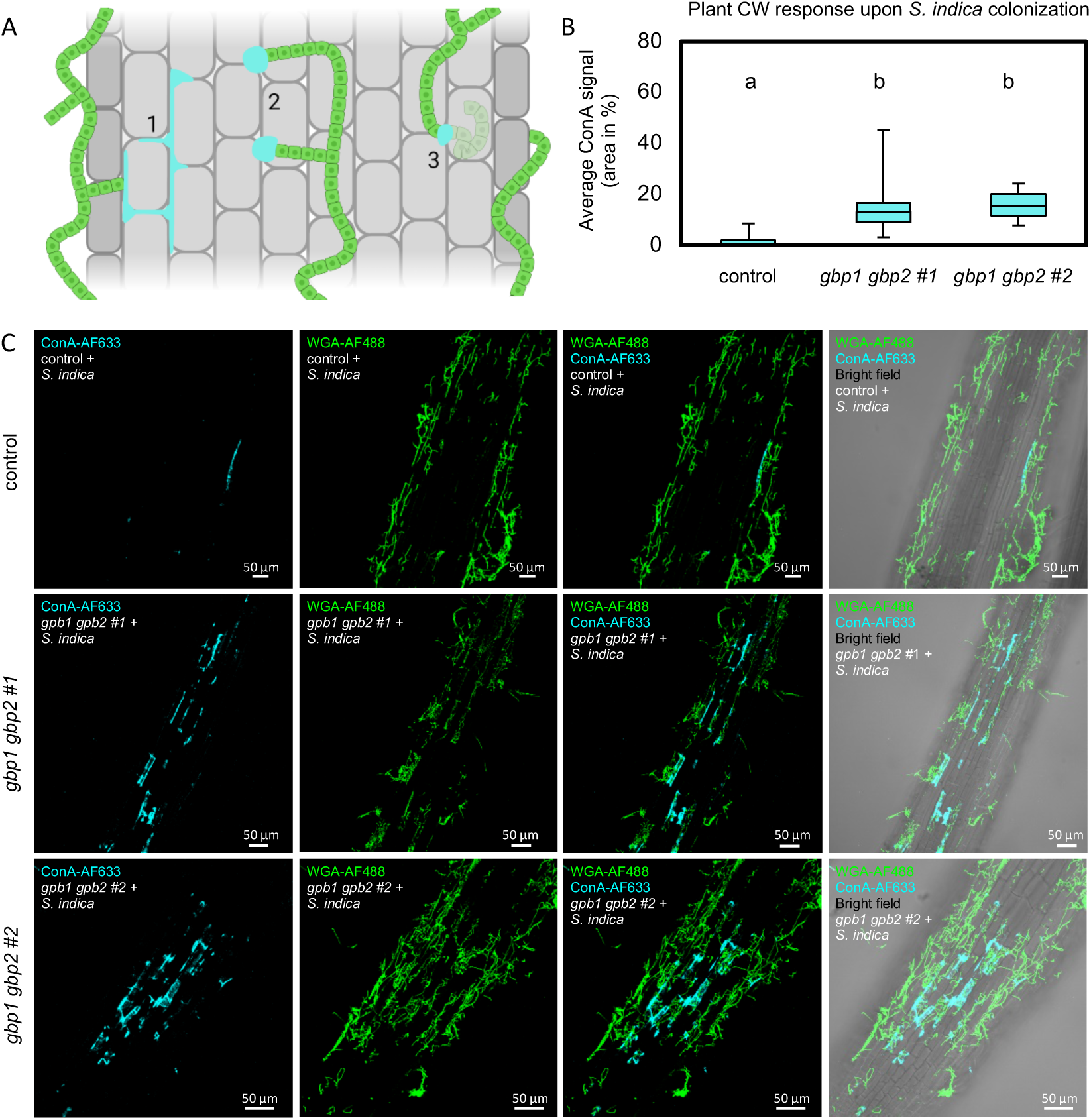
Colonization of barley *gbp1 gbp2* mutant lines by *S. indica* results in CW hyperresponse. (A) Schematic representation of different forms of CWAs in barley roots in response to fungal colonization, including an extensive reaction along the CWs (1), papillae formation at the sites of attempted (2) and successful (3) hyphal penetration. (B) Area of the root showing CW response in barley control and *gbp1 gbp2* mutant lines in percent. Letters represent statistically significant differences in ConA signal based on a Kruskal-Wallis test and Dunn’s post hoc test (significance threshold: P ≤0.05). (C) Barley root CW responses of control and *gbp1 gbp2* mutant lines colonized by *S. indica*. Samples were fluorescently labelled with concanavalin A (ConA-AF633, cyan) and wheat germ agglutinin (WGA-AF488, green) for visualization of CWAs and fungal structures, respectively, then analyzed by confocal laser scanning microscopy. ConA, concanavalin A, CW(A), cell wall (appositions), WGA, wheat germ agglutinin.

## Discussion

The success of fungal colonization is largely determined by the environmental conditions (e.g. nutrient availability) and genetic factors of the host. Depending on the type of plant-fungal interaction, these genetic factors are referred to as susceptibility factors (pathogenic interaction) or compatibility factors (beneficial interaction). Their mutation can interfere with fungal colonization at different stages by hampering tissue penetration and early establishment, by modulating the plant immune system or by removal of structural and metabolic components that are required for fungal maintenance (van Schie & Takken, 2014). In AM symbiosis, compatibility is mediated by a set of evolutionary conserved genes that are exclusively present in AM host species (Bravo *et al*., 2016; Radhakrishnan *et al*., 2020). In cooperation with membrane-integral LysM receptor-like kinases, these components are responsible for the signal transduction and decoding of nuclear calcium spiking signatures triggered by microbial chitin-derived oligomers. While this pathway is considered to be specific for the accommodation of AM fungi, many endophytes and pathogens have been shown to exploit it at least partially to ensure their own establishment within the host (Wang *et al*., 2012; Rey *et al*., 2013, 2015; Gobbato *et al*., 2013; Skiada *et al*., 2020). Many fungi have adapted their colonization strategies to fundamental developmental processes, for example by relying on the host’s CW structure and membrane dynamics for successful establishment within host tissue. Interferences of these processes can massively impact fungal compatibility (van Schie & Takken, 2014).

In this study, we present a previously undescribed role of GBPs, a family of GH81-type β-1,3-endoglucanases, as tissue-independent, broad-spectrum compatibility factors for fungal colonization. Barley GBP1 was identified as the major β-glucan interactor by a pull-down approach using laminarin as a β-1,3-glucan bait (Figure 1A). Laminarin is an algal storage carbohydrate which commonly serves as a commercially available surrogate substrate for fungal cell surface glycans due to its β-1,3/1,6-linkage pattern (Fesel & Zuccaro, 2016; Wanke *et al*., 2021). Plant β-1,3-glucanases have been known to be involved in plant development and plant-microbe interactions for a long time (Perrot *et al*., 2022). In the context of plant immunity, β-1,3-endoglucanases became known as pathogenesis-related group 2 (PR-2) proteins due to their crucial role in plant defense. Contrasting with the primary role of β-1,3-endoglucanases in plant resistance, we here show that knocking out the only two GBP paralogues in barley leads to reduced colonization by the mutualistic root endophytes *S. indica* and *S. vermifera* as well as the AM fungus *R. irregularis* (Figure 1). This suggests that barley GBPs serve as compatibility factors for colonization by beneficial root fungi. This compatibility principle could be extended beyond mutualistic fungi, as colonization by the hemibiotrophic root pathogen *B. sorokiniana* as well as the biotrophic foliar pathogen *B. graminis* was similarly hindered (Supplementary Figure 7). Fluorescence microscopy revealed that reduced colonization by the mutualistic fungus *S. indica* in the *gbp1 gbp2* mutants was accompanied by a strong response at the host CWs stained with the lectin ConA (Figure 4). These CWAs were not spatially restricted to the sites of fungal penetration but instead excessively and amorphously spread along the host CW within the colonized area (Figure 4). Since spontaneous CWA formation in mock-treated plants was not observed, constitutive activation of these defense responses by *gbp1 gbp2* mutation is unlikely (Supplementary Figure 14). The contribution of papillae formation to fungal resistance has been controversially discussed over the last decades, however, several studies reported that both early and increased papillae formation can protect hosts from fungal penetration (Hückelhoven, 2007, 2014). Arabidopsis plants constitutively expressing the callose synthase PMR4 produced enlarged callose depositions at the inner and outer side of the cellulosic CW at early time points of fungal infection (Ellinger *et al*., 2013; Eggert *et al*., 2014). This mediates complete penetration resistance to adapted and non-adapted powdery mildew strains without the induction of either SA- or JA- dependent pathways (Ellinger *et al*., 2013). Callose depositions induced by defense priming also enhanced resistance to the necrotrophic pathogens *Alternaria brassicicola* and *Plectosphaerella cucumerina* in Arabidopsis (Ton & Mauch-Mani, 2004). Moreover, *mlo (mildew resistance locus O)*-dependent resistance of barley to *B. graminis* and *S. indica* is accompanied by a faster formation of larger papillae in leaves or roots, respectively (Skou *et al*., 1984; Jørgensen, 1992; Chowdhury *et al*., 2014; Hilbert *et al*., 2019). Similar to our observations for the *gbp1 gbp2* mutants, barley *mlo* mutants also exhibited impaired colonization by AM fungi during early stages of interaction (Jacott *et al*., 2020). This decrease in colonization by mutualistic fungi may partially explain why genes contributing to pathogen infection have been conserved throughout evolution. Mutations of compatibility genes often bring along developmental penalties (e.g. growth, yield) or increased susceptibility to pathogens with other infection strategies (van Schie & Takken, 2014). However, we did not observe any differences in root and shoot fresh weight between the control lines and the *gbp1 gbp2* mutants (Supplementary Figure 5). While barley *mlo* mutants were more susceptible to necrotrophic and hemibiotrophic foliar pathogens due to the spontaneous occurrence of cell death (Jarosch *et al*., 1999; McGrann *et al*., 2014), barley *gbp1 gbp2* mutants exhibited reduced colonization by the aggressive hemibiotrophic root pathogen *B. sorokiniana*. Future studies should survey colonization rates of root (e.g. *Fusarium graminearum*, *Rhizoctonia solani*) and leaf (e.g. *Pyrenophora teres* f. sp*. teres*, *Rhynchosporium secalis*) necrotrophs.

The causal link between the *gbp1 gbp2* mutations and reduced fungal compatibility remains yet to be clarified. Barley GBP1 could process β-glucan-based host- or microbe-associated molecular patterns released during fungal colonization to modulate immune response activation as fungal colonization progresses. Missing GBP activity could lead to an overaccumulation of long-chain β-glucan elicitors in the host apoplast, possibly explaining the large-scale induction of CW responses observed in the *gbp1 gbp2* mutant background upon *S. indica* colonization. A related homeostasis principle was recently demonstrated for berberine bridge enzyme-like oxidases that counteract the deleterious accumulation of active oligogalacturonides and cellodextrins as DAMPs in the host apoplast during microbial colonization (Benedetti *et al*., 2018; Locci *et al*., 2019). GBPs could also be involved in the release or tailoring of a yet undescribed β-glucan-derived signaling molecule that dampens immunity to facilitate colonization of mutualistic fungi (Dumas-Gaudot *et al*., 1996). Similar to β-glucans, symbiotic signals such as short chitooligosaccharides (e.g. chitotetraose) and lipo-chitooligosaccharides are abundantly found in both symbiotic and pathogenic fungi and fulfill various functions relevant for inter-kingdom interactions (Feng *et al*., 2019; Rush *et al*., 2020). Alternatively, GBP1 could be an important factor for dynamic adjustment of CW responses. The β-1,3-glucan callose is - along with phenolic substances and other CW carbohydrates - a major component of CWAs (Chowdhury *et al*., 2014; Hückelhoven, 2014). GBPs could directly act on defense-triggered CWAs by degradation of their β-glucan components, thus acting as a negative regulator of CWA formation.

In the apoplastic space, the hydrolytic activity of β-1,3-endoglucanases contributes to host immunity in multiple ways (Perrot *et al*., 2022). Secreted β-1,3-glucanases directly process the β-glucan-containing surface glycans of invading fungi and oomycetes. Furthermore, this hydrolytic activity leads to the release of glucan fragments that can act as elicitors of plant defense responses. Since the outermost layer of fungal cell surface glycans mainly consists of β-1,3-glucans (Wanke *et al*., 2021), hydrolysis by β-1,3-glucanases renders the inner chitin layer more accessible to secreted plant chitinases. Although GH81 family members are not categorized as PR-2 proteins, previous studies focusing on GBPs from soybean and other legumes have shown their importance in defense against oomycetes (Fliegmann *et al*., 2004; Leclercq *et al*., 2008). To better understand the contribution of GBP1 to hydrolysis of fungal surface glycan substrates, we performed substrate characterization assays with heterologously purified barley GBP1. GBP1 specifically hydrolyzes linear β-1,3- glucans as short as laminaritriose (Figure 2C), but did not effectively act on preparations of the extracellular polysaccharide matrix or CW isolated from *S. indica* (Supplementary Figure 10). EPS matrices from a variety of plant-colonizing fungi consist of a β-1,3-linked glucan backbone heavily substituted with β-1,6-linked glucose (Wawra *et al*., 2019; Chandrasekar *et al*., 2022). Barley BGLUII, an enzyme secreted upon confrontation with fungi, has been shown to release a β-glucan decasaccharide with antioxidative properties from this EPS matrix that facilitates fungal colonization (Chandrasekar *et al*., 2022). Since GBP1 was unable to digest highly substituted laminarin from *E. bicyclis* (Pang *et al*., 2005; Mystkowska *et al*., 2018), we conclude that the high degree of branching in the EPS matrix similarly protects this cell surface glycan from hydrolysis by GBP1 (Supplementary Figure 11). However, we cannot exclude that cell surface glycans from other fungi or oomycetes might serve as better substrates for GBP1. Although GBP1 from barley did not exhibit a pioneering hydrolytic role on the tested preparations, in a more complex scenario with a variety of secreted host hydrolases, GBP1 could belong to a second set of hydrolytic enzymes acting on the predigested cell surface glycans of microbial invaders. In the performed ROS burst assays, we observed that treatment of the long β-glucan laminarin with GBP1 differentially modulated ROS production in the two tested species (Figure 3C and 3D), demonstrating that GBP1 has the potential to fine-tune ROS homeostasis.

Soybean GBPs contain two carbohydrate-associated domains: an endoglycosidic hydrolase domain and a high-affinity binding site for a *Phytophthora sojae*-derived β-glucan heptaglucoside elicitor (Fliegmann *et al*., 2004). Inhibition of elicitor binding to GBP with a specific antibody prevented the production of phytoalexins, suggesting that soybean GBP may be part of the β-glucan-heptaglucoside receptor complex (Umemoto *et al*., 1997). Since GBP1 was identified by a pull-down using laminarin, we hypothesize that GBP1 has a β-glucan binding domain in addition to the hydrolytic domain. However, barley *gbp1 gbp2* ROS production is not reduced upon treatment with various β-glucan elicitors, ruling out the long-standing assumption that barley GBPs are necessary components of a β-1,3-glucan receptor complex involved in immunity (Fliegmann *et al*., 2004). Functional differences between barley and legume GBPs are not surprising given the massive expansion of GBPs in the legume family, suggesting that legume GBPs may have diversified in terms of their role upon a multitude of various abiotic and biotic stresses and accommodation of beneficial microbes as suggested for the MLO family members (Jacott et al., 2020). Consistent with the facilitative role of barley BGLUII in fungal colonization, our findings provide new insights of how spatiotemporal dynamics in perception and hydrolysis of host and microbial β-glucans modulate fungal establishment in the host tissue.

For decades, plant β-1,3-glucanases were predominantly considered to be deployed as a defensive strategy to hinder microbial invasion. Our work shows that β-1,3- glucanases such as barley GBPs can play a previously undescribed role as compatibility factors for colonization by fungi of different phylogenetic origins and lifestyles. Our findings provide new insights into the contribution of host β-glucanases to spatiotemporal dynamics in perception and degradation of host and microbial β-glucans that promote fungal establishment in the host tissue.

## Material and methods

### Plant material and growth conditions

#### Hordeum vulgare

All experiments, including the generation of CRISPR/Cas9 knock-out lines, were performed with the spring barley (*H. vulgare* L.) Golden Promise Fast, an introgression line carrying the *Ppd-H1* allele that confers fast flowering (Gol *et al*., 2021). From here on, we use “control” to name this non-mutagenized cultivar that carries normal copies of *GBP1* and *GBP2*. For ROS burst assays, barley seeds were surface sterilized with 6% bleach for 1 hour and then washed extensively (5 × 30 mL sterile water). Seeds were germinated on wet filter paper at room temperature in the dark under sterile conditions for three days before transfer to sterile jars containing solid 1/10 plant nutrition medium (PNM), pH 5.7 and 0.4% Gelrite (Duchefa, Haarlem, the Netherlands) (Lahrmann *et al*., 2013). Seedlings were cultured for four days in a growth chamber under long-day conditions (day/night cycle of 16/8 h, 22°C/18°C, light intensity of 108 μmol·m^−2^·s^−1^).

#### Nicotiana benthamiana

Seeds of *N. benthamiana* wild-type lines were sown on soil and grown for 3 weeks in the greenhouse under long-day conditions (day/night cycle of 16/8 h, 22–25°C, light intensity of ∼140 μmol·m^−2^·s^−1^, maximal humidity of 60%).

### Carbohydrate substrates for immunity and enzymatic digestion assays

All laminarioligomeres, gentiobiose, chitohexaose, cellohexaose and xyloglucan oligomer (XXXG) were purchased from Megazyme (Bray, Ireland). Laminarin from *Laminaria digitata* was purchased from Sigma-Aldrich (Taufkirchen, Germany) and laminarin from *Eisenia bicyclis* was purchased from Biosynth (Staad, Switzerland). Substrates derived from *Serendipita indica* CW and EPS matrix were purified and prepared as previously described (Chandrasekar *et al*., 2022).

### Identification of protein candidates interacting with β-glucan

#### Biotinylation of laminarin

Laminarin (final concentration ∼ 60 mM) was incubated with biotin-hydrazide (120 mM) and sodium cyanoborohydride (1 M) for 2 h at 65°C. The product was purified on a PD MidiTrap G-10 column (GE Healthcare) according to the manufacturer’s description. Success of biotinylation was validated via mass spectrometry. The immunogenic capacity of biotinylated laminarin was confirmed via ROS burst assays.

#### Protein pull-down with biotinylated laminarin

Barley leaves from 2-week-old plants were treated for 15 min with biotinylated laminarin, followed by vacuum infiltration for 2 min. Untreated laminarin and a biotinylated version of the bacterial elongation factor Tu peptide (elf18) were used as controls. The tissue was frozen in liquid nitrogen and ground to fine powder. Then, 10 mg·ml^-1^ of extraction buffer (10mM MES, 50mM NaCl, 10mM MgCl2, 1mM DTT, 1% IGEPAL, proteinase inhibitor cocktail) were added to the powder. To avoid pH-dependent binding effects to the NeutrAvidin beads, two buffer conditions (pH 5.6 and pH 8.0) were used for all treatments. Samples were incubated rotating at 4°C for 60 min and centrifuged at 10,000 rpm, 4°C for 15 min. Supernatant was filtered to remove pieces, mixed with 50 µL of high-capacity Neutravidin agarose resin (Thermo Fisher Scientific, Schwerte, Germany), and incubated (inverting) at 4°C for 3 h. The sample was briefly centrifuged at 700 rpm for 1 min. After discarding the supernatant, the beads were washed four times with 10 mL of wash buffer (10mM MES [pH 5.6 or pH 8.0], 50mM NaCl,10mM MgCl2, 0.5% IGEPAL). Proteins were eluted by boiling the beads with 50-70 µL of 2 x SDS loading (including reducing agent) for 5 min. Proteins were separated by SDS-PAGE (NuPAGE; Invitrogen, Waltham, United States) after staining with Coomassie brilliant Blue G-250, cut out for mass spectrometric analysis and digested with trypsin.

#### Tandem mass spectrometric (MS-MS) analysis of the pull-down proteins

LC-MS/MS analysis was performed using an Orbitrap Fusion trihybrid mass spectrometer (Thermo Scientific) and a nanoflow-UHPLC system (Dionex Ultimate3000, Thermo Scientific). Peptides were trapped to a reverse phase trap column (Acclaim PepMap, C18 5 µm, 100 µm x 2 cm, Thermo Scientific) connected to an analytical column (Acclaim PepMap 100, C18 3 µm, 75 µm x 50 cm, Thermo Scientific). Peptides were eluted in a gradient of 3 - 40 % acetonitrile in 0.1 % formic (solvent B) acid over 86 min followed by gradient of 40-80 % B over 6 min at a flow rate of 200 nL·min at 40 °C. The mass spectrometer was operated in positive ion mode with nano-electrospray ion source with ID 0.02mm fused silica emitter (New Objective). Voltage +2200 V was applied via platinum wire held in PEEK T-shaped coupling union with transfer capillary temperature set to 275 °C. The Orbitrap, MS scan resolution of 120,000 at 400 m/z, range 300 to 1800 m/z was used, and automatic gain control (AGC) was set at 2e5 and maximum inject time to 50 ms. In the linear ion trap, MS/MS spectra were triggered with data dependent acquisition method using ‘top speed’ and ‘most intense ion’ settings. The selected precursor ions were fragmented sequentially in both the ion trap using CID and in the HCD cell. Dynamic exclusion was set to 15 sec. Charge state allowed between 2+ and 7+ charge states to be selected for MS/MS fragmentation. Peak lists in the format of Mascot generic files (mgf files) were prepared from raw data using MSConvert package (ProteoWizard). Peak lists were searched on Mascot server v.2.3 (Matrix Science) against an *Hordeum vulgare* Morex v1.0 database (IBSC_v2, IPK Gatersleben) and an in-house contaminants database. Tryptic peptides with up to two possible mis-cleavages and charge states +2, +3, +4, were allowed in the search. The following modifications were included in the search: oxidized methionine as variable modification and carbamidomethylated cysteine as static modification. Data were searched with a monoisotopic precursor and fragment ions mass tolerance 10 ppm and 0.6 Da, respectively. Mascot results were combined in Scaffold v. 4 (Proteome Software) and exported in Excel (Microsoft Office, Supplementary Table 1).

### Plasmid construction for the heterologous expression of barley GBP1 in N. benthamiana

For *in planta* protein production in *N. benthamiana*, we used the binary vector pXCScpmv-HAStrep (Witte *et al.,* 2004; Myrach *et al.,* 2017) characterized by a 35S promoter cassette, modified 5′- and 3′-UTRs of RNA-2 from the cowpea mosaic virus as translational enhancers, and C-terminal hemagglutinin (HA) and StrepII tags. The codon-optimized GBP1 coding sequence was amplified with the primer pair *ClaI*_GBP1_F (5’-gacggtatcgataaaATGCCGCCACATGGTAGACG-3’) and GBP1_noSTOP_*XmaI*_R (5’-ataactcccgggATGGCCATATTGACGATACCAACAGC- 3’) and directionally cloned into the *ClaI* and *XmaI* sites of the binary vector to produce pXCScpmv-GBP1-HAStrep. To generate a catalytically inactive version of GBP1, the first glutamate residue of the catalytic center (E500) was exchanged to an alanine residue via site-directed mutagenesis PCR with the primer pair GBP1_E500A_F (5’- CAGG<u>C</u>ATCAACATCAGAAGCAGTG-3’) and GBP1_E500A_R (5’-GTTCCTACCATCTCCAAACTCAGTC-3’). The linearized, mutated plasmid was purified after gel electrophoresis using the NucleoSpin Gel and PCR Clean-up Kit (Machery-Nagel, Düren, Germany). The isolated DNA fragment was treated with a self-made KLD mixture (1,000 units·ml^-1^ T4 polynucleotide kinase, 40,000 T4 DNA ligase units·ml^-1^ ligase 2,000 units·ml^-1^ DpnI, 1 × T4 DNA ligase buffer; all enzymes were purchased from New England BioLabs, Ipswich, USA) for 1 hour at room temperature before transformation into *Escherichia coli* MachI cells. Plasmids were isolated using the NucleoSpin Plasmid Kit (Machery-Nagel, Düren, Germany) and sequenced to confirm the introduced mutation. Both plasmids (pXCScpmv-GBP1- HAStrep and pXCScpmv-GBP1_E500A-HAStrep) were introduced into *Agrobacterium tumefaciens* GV3101::pMP90RK strains for transient transformation of *N. benthamiana* leaf tissue.

### Heterologous protein production and purification from N. benthamiana

*A. tumefaciens* GV3101::pMP90RK strains carrying the binary vectors for protein production (antibiotic selection: 30 µg·mL^-1^ Rifampicin, 25 µg·mL^-1^ Kanamycin, 50 µg·mL^-1^ Carbenicillin) and *A. tumefaciens* GV3101 strains carrying the binary vector for viral p19 silencing inhibitor expression (antibiotic selection: 30 µg·mL^-1^ Rifampicin, 30 µg·mL^-1^ Gentamicin, 100 µg·mL^-1^ Carbenicillin) were grown in selection LB liquid medium at 28°C, 180 rpm for three days. The cultures were centrifuged (3,500 g for 15 min), resuspended in infiltration buffer (10 mM MES pH 5.5, 10 mM MgCl2, 200 μM acetosyringone) to an OD600 of 1 and incubated for 1 h in the dark at 28°C, 180 rpm. Each of the two *A. tumefaciens* strains carrying the GBP1 production constructs was mixed with the *A. tumefaciens* strain carrying the p19-expressing construct in a 1:1 ratio. The bacterial suspensions were infiltrated into the four youngest, fully developed leaves of four-week-old *N. benthamiana* plants with a needleless syringe. Five days after infiltration, the leaves were detached from the plant and ground in liquid nitrogen. Protein purification was carried out according to Werner *et al*. (2008) with minor modifications: The ground plant material (up to the 5 mL mark of 15-mL tube) was thoroughly resuspended in 5 mL of ice-cold extraction buffer (100 mM Tris pH 8.0, 100 mM NaCl, 5 mM EDTA, 0.5% Triton X-100, 10 mM DTT, 100 μg·ml^-1^ Avidin) and centrifuged at 10,000 g, 4°C for 10 min. The supernatant was filtered through a PD-10 desalting column (Sigma-Aldrich, Taufkirchen, Germany), transferred to a new tube, and supplemented with 75 μl·ml^-1^ Strep-Tactin Macroprep (50% slurry) (IBA Lifesciences GmbH, Göttingen, Germany). Samples were incubated in a rotary wheel at 4°C for one hour, followed by centrifugation for 30 s at 700 g. The supernatant was discarded, and the beads were washed three times with 2 mL of washing buffer (50 mM Tris pH 8.0, 100 mM NaCl, 0.5 mM EDTA, 0.005% Triton X-100, 2 mM DTT). Proteins were eluted from the beads by adding 100 μL of elution buffer (wash buffer containing 10 mM biotin) and incubating at 800 rpm for 5 min at 25 °C. The samples were centrifuged at 700 g for 20 s and the elution was repeated two more times. The elution fractions were pooled and dialyzed overnight against cold MilliQ water (dialysis tubing with 6-8 kDa cut-off). Proteins were stored on ice at 4°C for further use. The success of protein purification was analyzed by SDS PAGE and Western Blotting (Werner *et al*., 2008).

### Oxidative burst assay

#### Preparation of the plant material

For immunity assays in barley, the roots and shoots of seven-day-old seedlings were separated. The root tissue between 2 cm below the seed and 1 cm above the tip was cut into 5 mm pieces. Each assay was carried out with randomly selected root pieces from 16 barley seedlings. Four root pieces were transferred to each well of a 96-well microtiter plate containing 150 µL of sterile Milli-Q water. Barley shoot assays were performed on 3-mm leaf discs punched from the youngest leaves of eight individual barley seedlings.

For immunity assays in *N. benthamiana*, 3-mm leaf discs from the youngest, fully developed leaf of eight three-week-old plants were transferred to a 96-well plate filled with 150 µL of sterile Milli-Q water.

#### Assay protocol

The ROS burst assay was based on previously published protocols (Chandrasekar *et al*., 2022; Felix *et al*., 1999). In brief, a 96-well plate containing water and plant material (as described above) was incubated overnight at RT to remove unstable contaminants that had resulted from mechanical damage to the tissue during preparation (e.g. ROS). The next day, the water was replaced with 100 µL of fresh Milli-Q water containing 20 µg·mL^-1^ horseradish peroxidase (Sigma-Aldrich, Taufkirchen, Germany) and 20 µM L- 012 (Wako Chemicals, Neuss, Germany). After a short incubation period (∼15 min), 100 µL of double-concentrated elicitor solutions were added to the wells. All elicitors were dissolved in Milli-Q water without additional treatment. Measurements of elicitor-triggered apoplastic ROS production were started immediately and performed continuously with an integration time of 450 ms in a TECAN SPARK 10 M multiwell plate reader (Männedorf, Switzerland).

### Enzymatic carbohydrate digestion and thin layer chromatography (TLC)

Carbohydrate digestion assays were performed using either purified barley GBP (heterologously expressed in *N. benthamiana*) or barley BGLUII (Chandrasekar *et al*., 2022; available from Megazyme, E-LAMHV). Preparations of the fungal CW and EPS matrix were incubated overnight in sterile MilliQ water at 65°C prior to enzymatic digestion. Substrate and enzyme concentrations, buffer compositions, digestion temperature and time are described in the figure captions. Digestion was stopped by denaturing the enzymes at 95°C for 10 min and the digestion products were stored at -20 °C prior to use. An aliquot of each sample was subjected to TLC using a silica gel 60 F254 aluminum TLC plate (Merck Millipore, Burlington, USA), using a running buffer containing ethyl acetate/acetic acid/methanol/formic acid/water at a ratio of 80:40:10:10:10 (v/v). D-glucose, laminaribiose β-1-3-(Glc)2, laminaritriose β-1-3- (Glc)3, gentiobiose β-1-6-(Glc)2, and laminaripentaose β-1-3-(Glc)5 at a concentration of 1.5 mg·mL^−1^ were used as standards (Megazyme, Bray, Ireland). To visualize the glucan fragments, the TLC plate was sprayed with glucan developer solution (45 mg N-naphthol, 4.8 mL H2SO4, 37.2 mL ethanol and 3 mL water) and baked at 95°C until the glucan bands became visible (approximately 4-5 min).

### MALDI-TOF analysis

The digested products of GBP1 were analyzed using Oligosaccharide Mass Profiling as previously described (Günl *et al*., 2011). Briefly, 2 µL of the samples were spotted onto 2 µL of crystalized dihydroxy benzoic acid matrix (10 mg mL^−1^) and analyzed by MALDI-TOF mass spectrometry (Bruker rapifleX instrument, Bremen). Mass spectra were recorded in linear positive reflectron mode with an accelerating voltage of 20,000 V. The spectra of the samples were analyzed using the flexanalysis software 4.0 (Bruker Daltonics, Billerica, USA).

### CRISPR/Cas9-based mutagenesis of GBP homologues in barley

The CRISPOR web tool (version 4.97, Concordet and Haeussler, 2018) was used to design two single guide RNAs for each *GPB1* and *GBP2*:

GBP1_gRNA1: 5’-CCCGGCACGCTTCTTCGCGC<u>CGG</u>-3’

GBP1_gRNA2: 5’-TGGCGCCTTCGGATGAACAG<u>CGG</u>-3’

GBP2_gRNA1: 5’-TAAGATCCGTCGAGGCAGTA<u>TGG</u>-3’

GBP2_gRNA2: 5’-GTACAGCCGTTGCTACCCGA<u>CGG</u>-3’

Golden Gate cloning was used to load each guide sequence from complementary oligonucleotides into shuttle vectors pMGE625, pMGE626, pMGE628, pMGE629. The four guide expression cassettes were then assembled into binary vector pMGE599 to create pMP202. Vectors and cloning protocols have been previously described (Kumar *et al*., 2018) and were kindly provided by Johannes Stuttmann. Stable transformation of pMP202 in Golden Promise Fast was performed as described (Amanda *et al*., 2022).

### Fungal colonization assays

#### Serendipita vermifera, Serendipita indica and Bipolaris sorokiniana

The growth conditions for barley, *S. vermifera* (MAFF305830, *Sv*)*, S. indica* (DSM11827, *Si*) and *B. sorokiniana* (ND90Pr, *Bs*) and the preparation of fungal suspensions for plant inoculation have been described previously (Sarkar *et al*., 2019; Mahdi *et al*., 2021). Four-day-old barley seedlings were transferred to sterile jars on 1/10 PNM (pH 5.7) and inoculated with 3 mL of either sterile water as control, *Sv* mycelium (1 g per 50 mL), *Bs* conidia (5,000 spores·mL^-1^) or *Si* chlamydospores (500,000 spores·mL^-1^). Plants were grown on a day/night cycle of 16/8 h at 22/18 °C and 60 % humidity under a light intensity of 108 μmol·m^−2^·s^−1^. Plant roots were harvested six days after inoculation (dpi), washed thoroughly to remove extraradical fungal hyphae, and frozen in liquid nitrogen. Four barley plants were pooled per biological replicate. RNA extraction to quantify endophytic fungal colonization, cDNA generation and RT-PCR were performed as previously described (Sarkar *et al*., 2019). The primers used are listed in Supplementary Table 2.

#### Blumeria graminis f. sp. hordei

Control and *gbp* knock-out lines were grown in soil in a growth chamber (polyclimate) at 19 °C, 60 % relative humidity and a photoperiod of 14 h with a light intensity of 100 μmol m^-2^ s^-1^. Primary leaves of seven-day-old plants were cut and placed with the adaxial side down on 1% plant agar plates before gravity inoculation with *B. graminis* f. sp. *hordei* isolate K1 at a conidial density of about 20 conidia per mm. For inoculation, ten-to fourteen-day-old plants were used for inoculation. Evaluation of disease symptoms was performed with five to seven days infected plants.

Leaves were harvested at the indicated time points and cleared in 70 % ethanol before staining of the fungal structures with Coomassie brilliant blue solution (0.1 % [w/v] Coomassie Brilliant Blue R250 in 50 % ethanol and 10 % acetic acid) for 10 to 15 seconds. Bright field microscopy was used to assess secondary hyphae formation in 50 germinated conidia spores in the tip area and 50 germinated conidia in the middle of the leaf. At least six independent leaves were examined at 48 h after infection for fungal penetration success.

#### Rhizophagus irregularis

Barley control and mutant seeds were sterilized and germinated as described previously (Mahdi *et al*., 2021). Germinated seedlings were transferred to 9 × 9 cm pots containing autoclaved 1:1 silica sand:vermiculite mixture inoculated with 700 mg *R. irregularis* spore inoculum (10,000 spores·g^-1^ moist diatomaceous earth powder [50% water]) (Symplanta, Darmstadt, Germany). Approximately 500 mg of the inoculum was evenly mixed into the lower two-third substrate layer, further 200 mg were evenly sprinkled into a hole in the upper third of the substrate layer, into which the seedlings were transplanted. The seedlings were grown in the greenhouse and watered weekly with 30 mL of deionized water and tap water in a 1:1 ratio. Roots were harvested at 28 dpi and stored in 50% EtOH at 4°C until staining.

Roots were stained according to a previously published protocol (Vierheilig *et al*., 1998). Briefly, roots were incubated for 15 min at 95°C in 10% KOH, washed with 10% acetic acid and incubated for 5 min at 95°C with a staining solution of 5% ink (Pelikan, Falkensee, Germany) in 5% acetic acid. After staining, the roots were carefully washed with tap water, then incubated in 5% acetic acid at 4°C for at least 20 minutes. The ink-stained root tissue was cut into 1 cm segments with a scalpel and 30 segments of similar diameter were randomly selected from each genotype. Cross-section points were determined from 10 random cuts per root segment. Ink-stained *R. irregularis* structures such as intraradical hyphae (IRH), extraradical hyphae (ERH), arbuscules and vesicles were visualized with a light microscope (AxioStar, Carl Zeiss, Jena, Germany) at 10× magnification. Colonization with *R. irregularis* was scored as positive if IRH, arbuscules or vesicles were present. The roots of four biological replicates per genotype were examined.

### Staining for confocal microscopy

Root tissue of barley control and mutant plants colonized by *S. indica* was harvested at 6 dpi and then stained as previously described (Hilbert *et al*., 2019). Briefly, roots were incubated at 95 °C for 2 min in 10% KOH, washed 3 times for 30 min in deionized water and 3 times for 30 min in PBS (pH 7.4). The roots were stained for 5 min under vacuum and then washed three times with deionized water. Fungal structures were visualized using 10 μg·mL^-1^ fluorescently labeled wheat germ agglutinin (WGA-AF488, Invitrogen, Thermo Fisher Scientific, Schwerte, Germany) in PBS (pH 7.4) and imaging was conducted with an excitation wavelength of 488 nm and emission detection between 500-540 nm. Papillae and root cell wall appositions were stained with 10 μg·mL^-1^ fluorescently labeled Concanavalin A (ConA-AF633, Invitrogen, Thermo Fisher Scientific, Schwerte, Germany) in PBS (pH 7.4) and imaged by excitation at 633 nm and detection at 650-690 nm.

Images were taken with a Leica TCS SP8 confocal microscope (Wetzlar, Germany). The percentage area of ConA staining was quantified using Fiji (Schneider *et al*., 2012) in maximum intensity projections of 10-slice Z-stacks with an image depth of 10 μm. At least 24 different root regions of each genotype were analyzed.

### Statistical analyses

A summary of the statistical analyses can be found in Supplementary Table 3.

### Data availability

Proteomic data described in this study has been deposited into the ProteomeXchage Consortium via PRIDE (Perez-Riverol *et al.,* 2019) partner repository with the dataset identifier: PXD0xxxxx and 10.xxxx/PXD0xxxxx.

### Accession numbers

*Hordeum vulgare* GBP1: HORVU5Hr1G059430.56 (*Hordeum vulgare* Morex v1.0, IBSC_v2, IPK Gatersleben),

*Hordeum vulgare* GBP2: HORVU6Hr1G034610.3 (*Hordeum vulgare* Morex v1.0, IBSC_v2, IPK Gatersleben),

*Hordeum vulgare* BGLU2: P15737 (UniProt), HORVU3Hr1G105630.7 (*Hordeum vulgare* Morex v1.0, IBSC_v2, IPK Gatersleben).

## Acknowledgements

We thank Hanna Rovenich, Dennis Mahr, Daniela Niedeggen and Edelgard Wendeler for technical support. We thank Gregor Langen for bioinformatic support and for discussions. AZ, MP, PS, IMLS and BC acknowledge support from the Cluster of Excellence on Plant Sciences (CEPLAS) funded by the Deutsche Forschungsgemeinschaft (DFG, German Research Foundation) under Germany’s Excellence Strategy–EXC 2048/1–Project ID: 390686111. AZ and LM acknowledge support from project ZU 263/11-1 (SPP 2125 DECRyPT). AW, LKM and SvB acknowledge support by the International Max Planck Research School (IMPRS) on ‘Understanding Complex Plant Traits using Computational and Evolutionary Approaches’ and the University of Cologne. IMLS was funded by the DFG Emmy Noether Programme (SA 4093/1-1). PD, NH, FLHM and CZ acknowledge support from The Gatsby Charitable Foundation.

## Author contributions

AW, SW and AZ conceived the study. SW, NH, PD, FLHM and CZ directed and supervised the protein pull down. IFA and AW generated the barley CRISPR/Cas9 mutants. LKM, SvB, MB and IMLS performed the fungal colonization assays. MN, PS and AW performed the biochemical experiments. BC, AZ and MP directed and supervised the carbohydrate analytics. AW and AZ supervised the project and designed the experiments. AW, SvB and AZ wrote the paper with contribution of all the authors.

