## Supplementary Figures for "A GH81-type β-glucan-binding protein facilitates colonization by mutualistic fungi in barley"

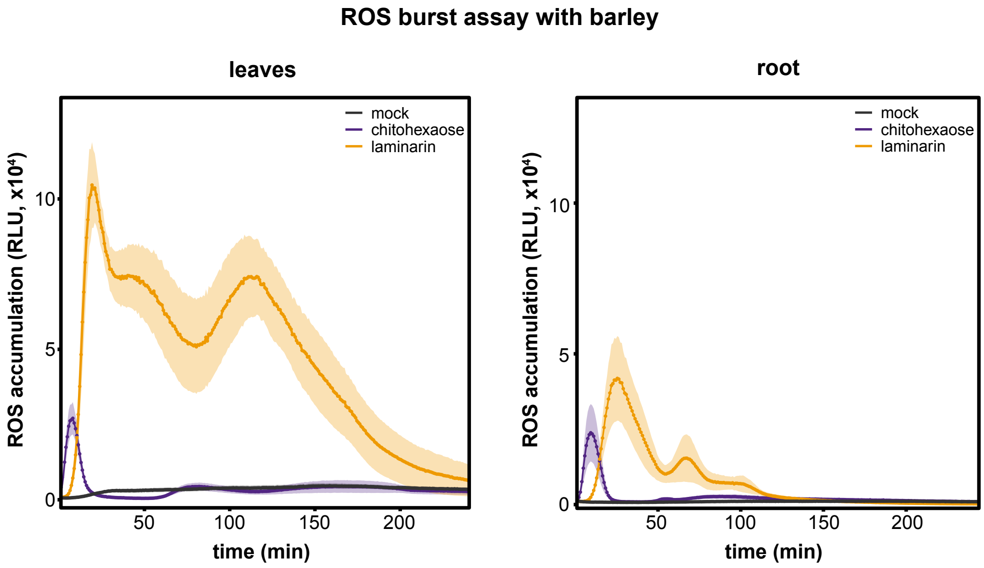


**Supplementary Figure 1. Barley leaf and root tissues respond similarly to laminarin treatment.** Apoplastic ROS accumulation after treatment of seven-day-old barley root pieces and leaf discs with chitohexaose (10 µM) and laminarin (4 mg·mL$^{-1}$). Treatment with Milli-Q water was used as mock control. Values represent mean ± SEM from 16 wells. The experiment was repeated at least three times with similar results. ROS, reactive oxygen species; RLU, relative luminescence units.

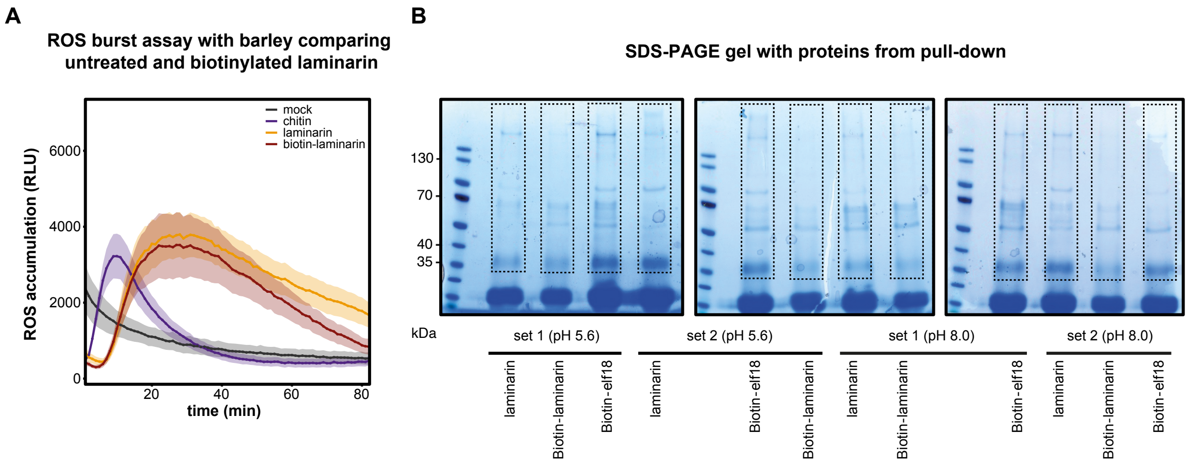


**Supplementary Figure 2. Protein pull-down with biotinylated laminarin in barley leaves.** (A) Apoplastic ROS accumulation after treatment of two-week-old barley leaf discs with untreated and biotinylated laminarin (each 6 mg·mL$^{-1}$). Treatment with Milli-Q water was used as mock control. Values represent mean ± SEM from 8 wells. (B) Separation of pull-down samples on SDS-PAGE gel prior to submission for mass spectroscopy. Pull-down was performed with untreated laminarin, biotinylated laminarin and biotinylated elf18 at two different pH values. Areas indicated by dotted lines were excised from the gel and further processed for mass spectrometric analyses. Elf18, peptide from bacterial elongation factor Tu; ROS, reactive oxygen species; RLU, relative luminescence units.

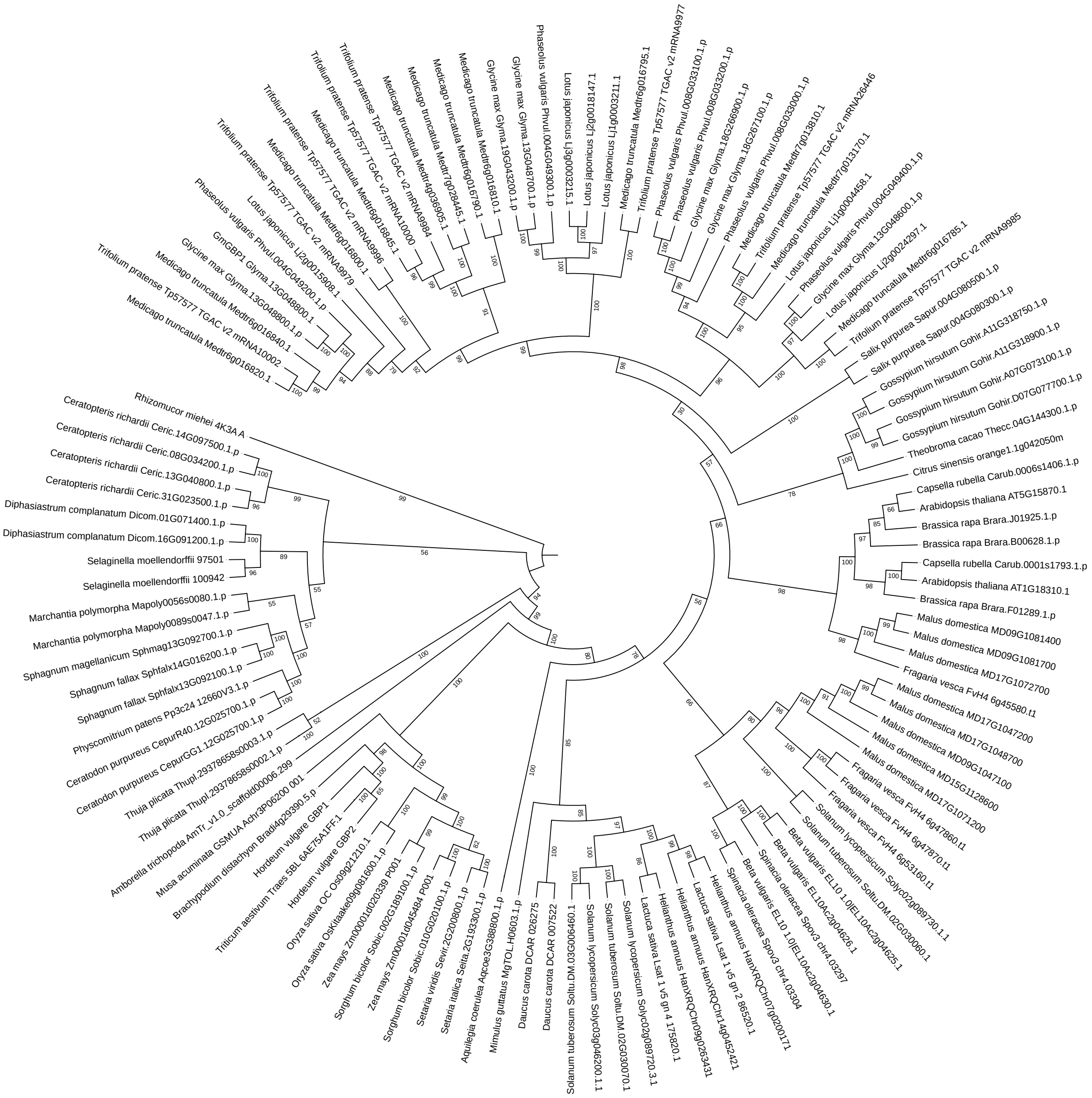


**Supplementary Figure 3. Duplication and expansion of GBP orthologues occurred in various plant lineages.** Homologues of barley GBP1 and GBP were obtained from the Phytozome database (min. half of query protein sequence covered, e-value≤0.001). The sequences were aligned with MAFFT (version 7.515), the alignment was trimmed with trimal (version 1.4, *automated1* setting) and a phylogenetic tree was constructed with IQ-TREE 2.2.0.3 (version, combined ModelFinder, tree search, SH-aLRT test with 10000 repetitions). Visualization of phylogenetic tree was performed with iTOL online software.

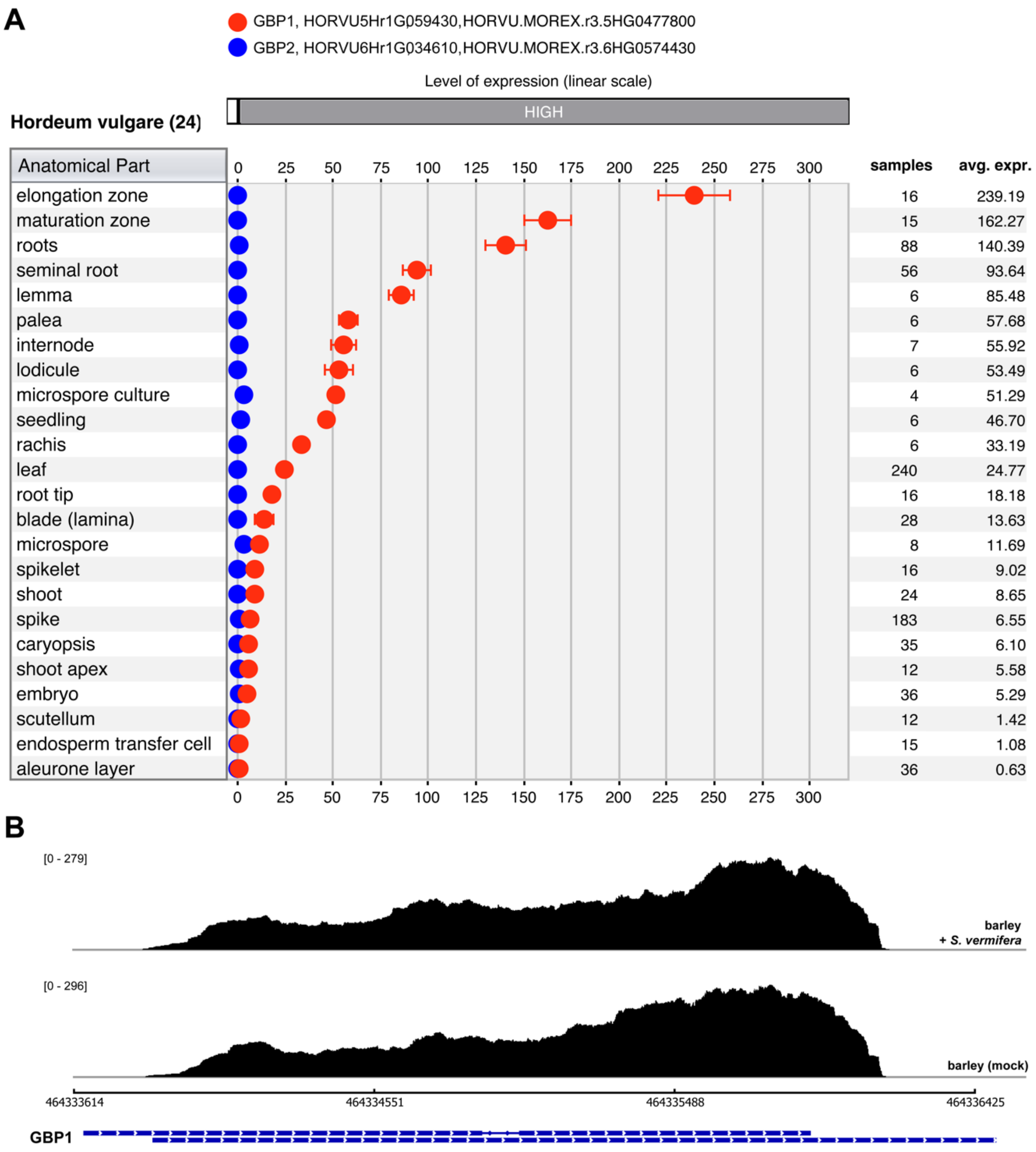


**Supplementary Figure 4. *GBP1* but not *GBP2* is expressed in different barley tissues.** (A) Figure based on expression datasets created by GENEVESTIGATOR. (B) Sashimi plots for *GBP1* using aligned reads from an RNA-seq experiment with barley roots colonized by *S. vermifera* (6 dpi) or corresponding non-colonized control (mock). No reads mapped to *GBP2* gene. Plots were generated with the Integrative Genome Viewer based on previously published data (Sakar *et al.,* 2019).

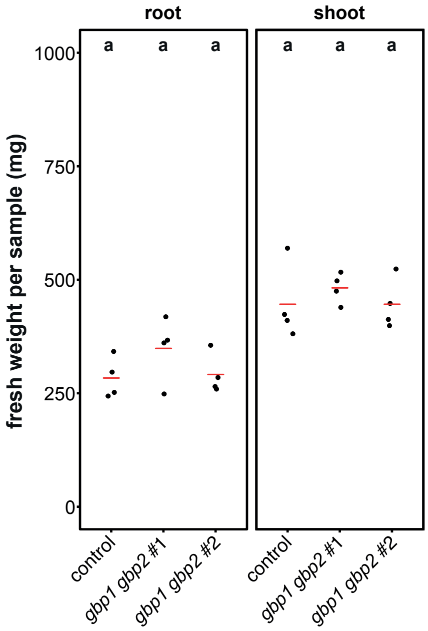


**Supplementary Figure 5. Fresh weight of root and shoot tissue of the barley control and *gbp1 gbp2* mutant lines.** The control line and *gbp1 gbp2* mutant lines were germinated on wet filter paper for 4 days and then grown on 1/10 PNM medium for 6 days under sterile conditions. Root and shoot fresh weight of the 10-day-old barley plants were measured. Black dots represent biological replicates and the red bar indicates the average fresh weight (n = 4, each replicate consists of 4 barley plants). Different letters represent statistically significant differences based on one-way ANOVA and Tukey's post hoc test using a 95% confidence interval.

**A**

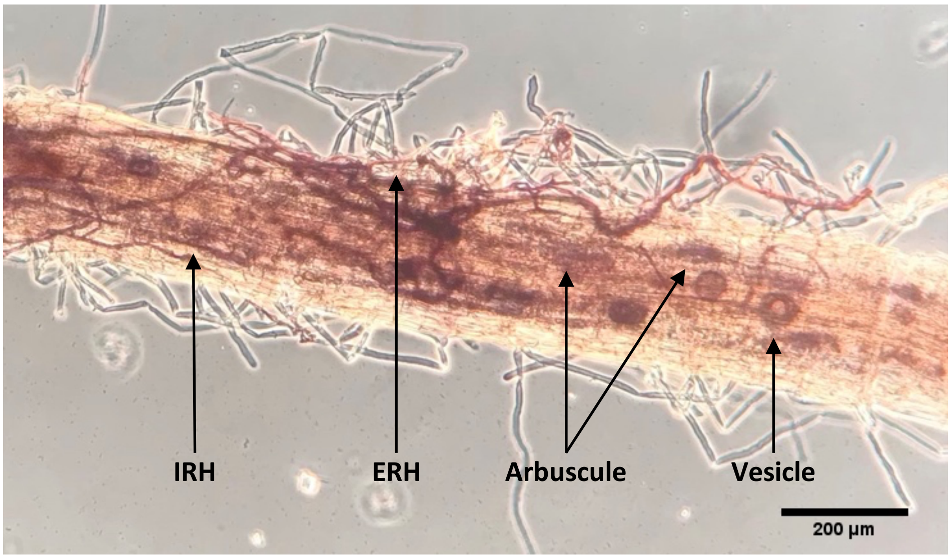


**B**

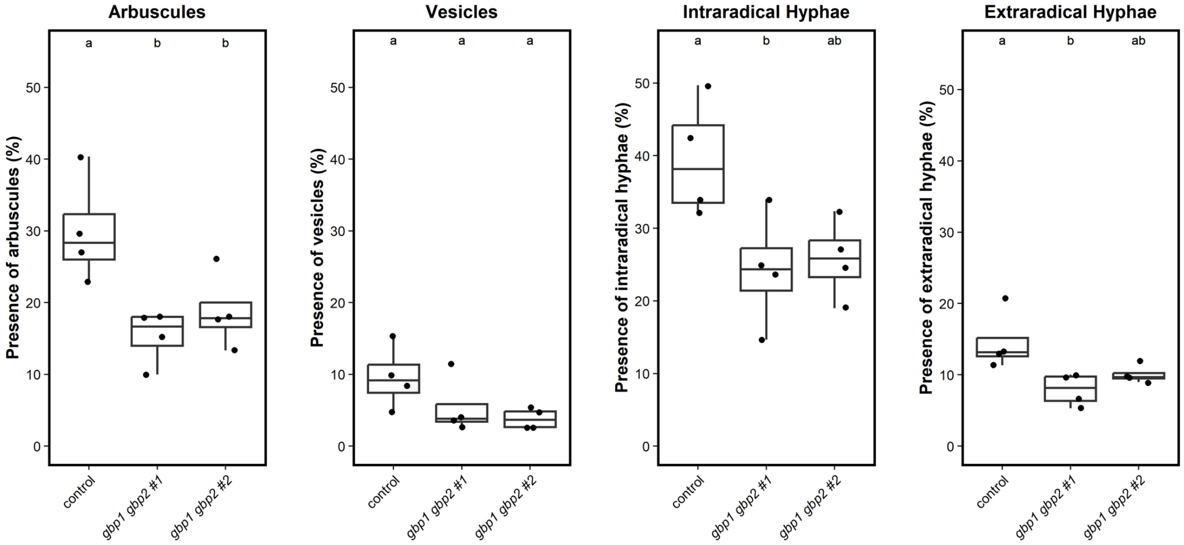


**Supplementary Figure 6. *R. irregularis* structures observed during colonization of the barley control and *gbp1 gbp2* mutant lines.** (A) Example of *R. irregularis* structures in barley roots at 28 dpi. *R. irregularis* structures were stained with 5% ink (Pelican) and fungal structures were observed using a light microscope (AxioStar, Carl Zeiss, Jena, Germany) at 10X magnification. (B) The control line and *gbp1 gbp2* mutants were inoculated with *R. irregularis* spores and roots were harvested at 28 dpi. Barley roots were analyzed at 300 randomly chosen sections (covering 30 cm of root length) for the presence of *R. irregularis* structures including arbuscules, vesicles, intraradical hyphae (IRH), and extraradical hyphae (ERH). All data points are plotted on graphs as circles (n=4). Boxplot elements in this figure: center line, median; box limits, upper and lower quartiles; whiskers, 1.5 × interquartile range. Different letters represent statistically significant differences based on one-way ANOVA and Tukey's post hoc test with significance threshold: $P \leq 0.05$.

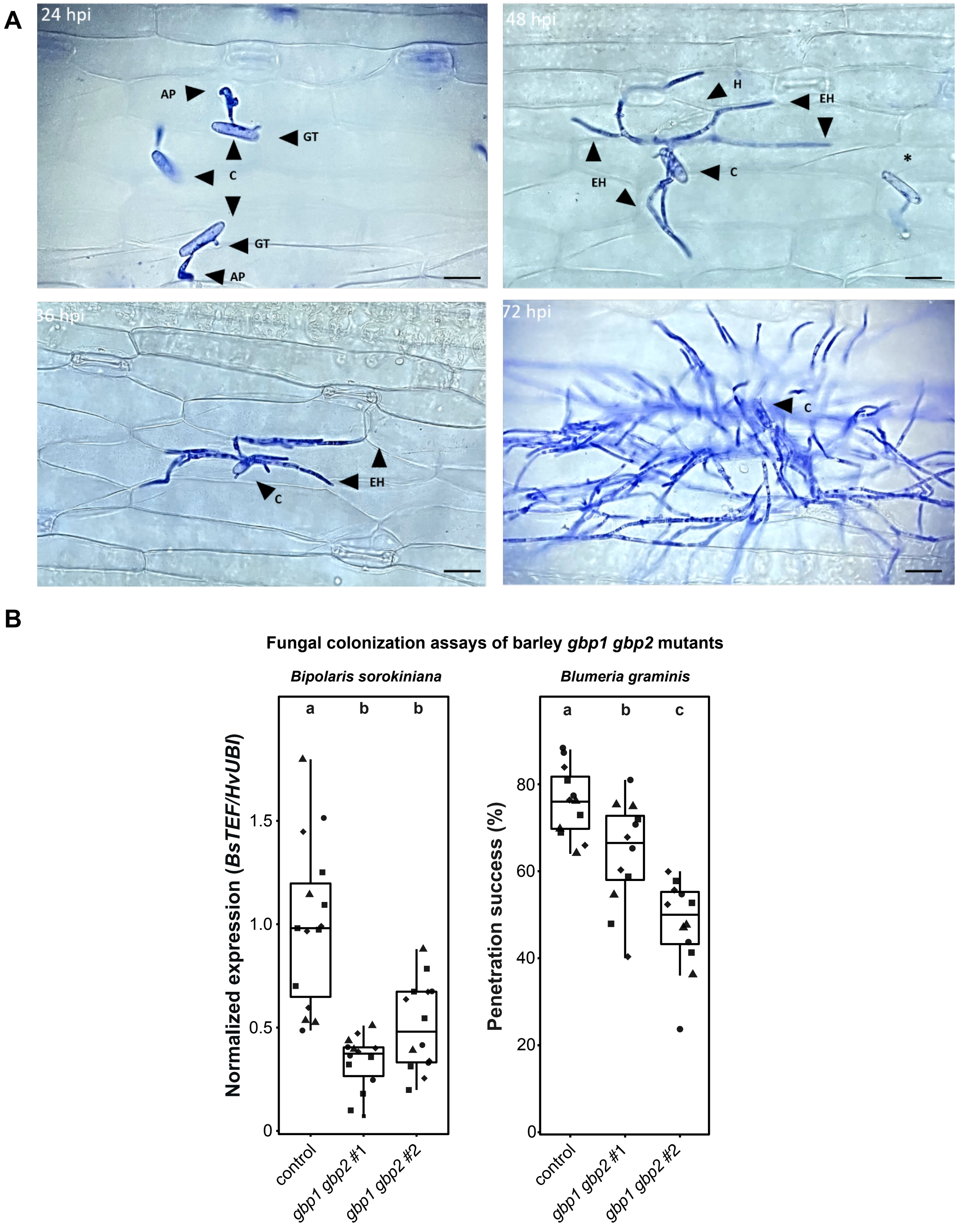


**Supplementary Figure 7. Colonization of barley control and *gbp1 gbp2* mutant lines by *B. sorokiniana and B. graminis* f.sp. *hordei.*** (A) Representative pictures of *B. graminis f.sp. hordei* infection structures on barley leaves. Barley leaves colonized by *B. graminis* were analyzed for penetration success using bright field microscopy. Fungal structures were stained using Coomassie brilliant blue, and secondary hyphae formation was assessed from 50 germinated conidia spores in both the tip and middle area of each leaf. (B) In barley root tissue, the expression of *B. sorokiniana* housekeeping gene *BsTEF* was quantified by RT-PCR and normalized to the barley housekeeping gene ubiquitin *HvUBI* (left graph). Quantification of *B. graminis* penetration success at 48 h (right graph). Boxplot elements in this figure: center line, median; box limits, upper and lower quartiles; whiskers, 1.5 × interquartile range. Data points from independent experiments are indicated by data point shape. AP, appressorium; C, conidia; EH, elongated secondary hyphae; GT, germ tube; H, hyphae. Different letters represent statistically significant differences based on one-way ANOVA and Tukey's post hoc test with significance threshold: $P \leq 0.05$.

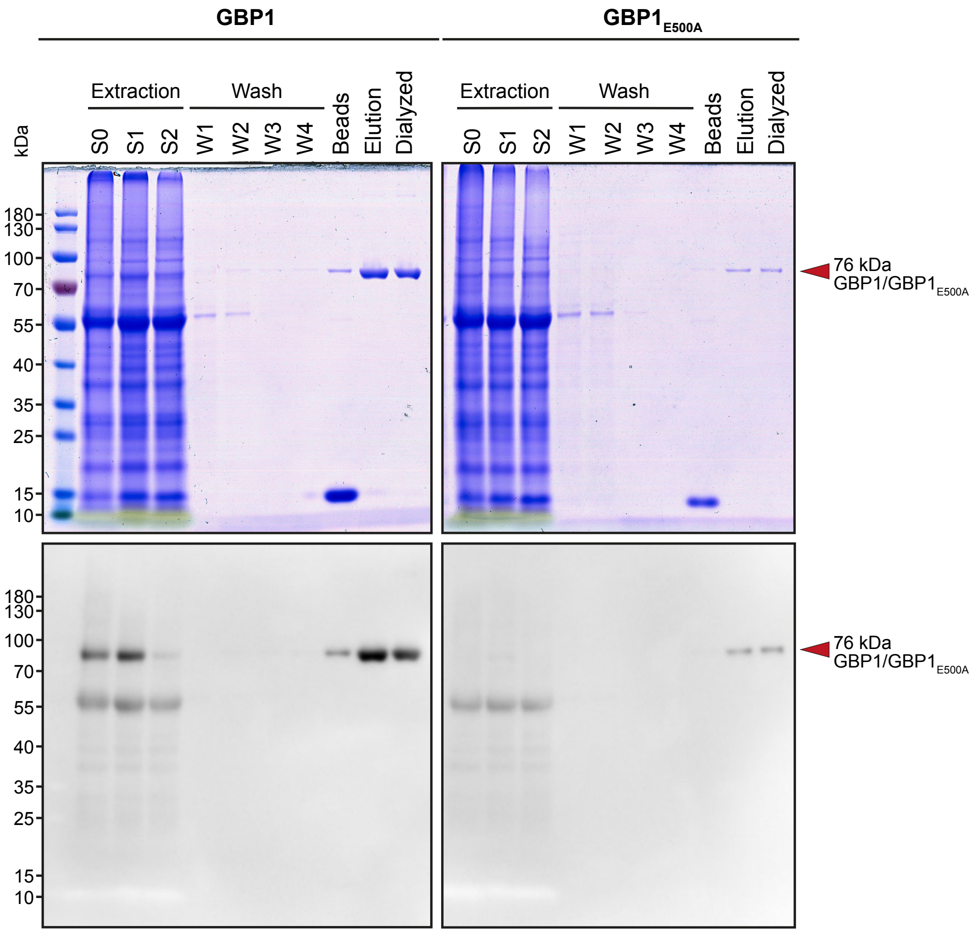


**Supplementary Figure 8. Purification of heterologously produced wildtype GBP1 and the mutated version GBP1$_{E500A}$ from *N. benthamiana* leaves.** Proteins were heterologously produced by Agrobacterium-mediated transient transformation and purified from *N. benthamiana* leaves using Strep-tag-II-based affinity chromatography. Protein purification success was analyzed via SDS-PAGE (upper row) and western blotting using an HRP-conjugated Strep-tag-II antibody (lower row). S0, crude sample; S1, sample after initial centrifugation; S2, sample after passing desalting column; W1-W4, washing steps; beads, Strep-Tactin beads after elution; elution, eluted fraction; dialyzed, eluted fraction after dialysis.

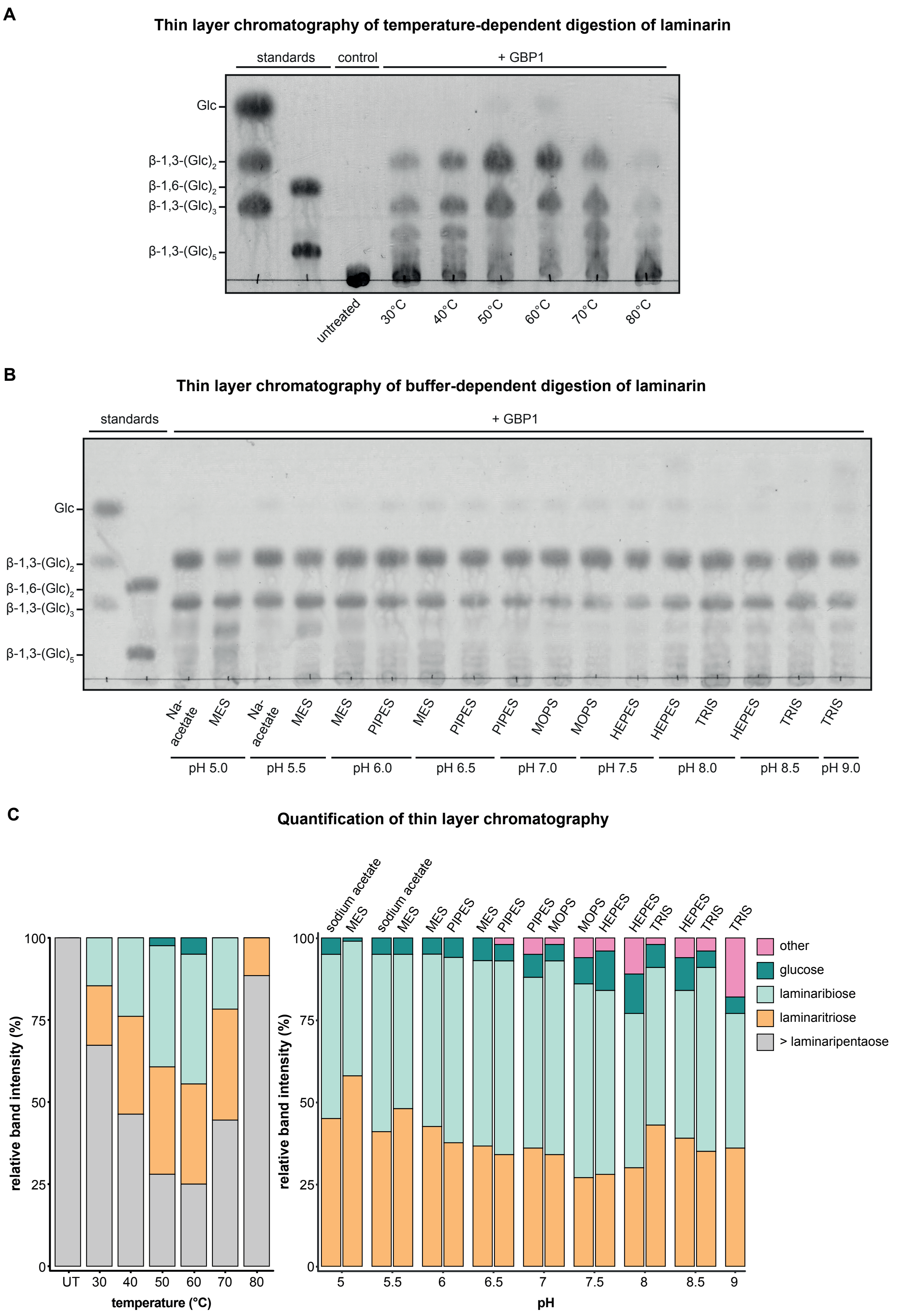
A
Thin layer chromatography of temperature-dependent digestion of laminarin
standards
control
+ GBP1
Glc
β-1,3-(Glc)2
β-1,6-(Glc)2
β-1,3-(Glc)3
β-1,3-(Glc)5
untreated
30°C
40°C
50°C
60°C
70°C
80°C
B
Thin layer chromatography of buffer-dependent digestion of laminarin
standards
+ GBP1
Glc
β-1,3-(Glc)2
β-1,6-(Glc)2
β-1,3-(Glc)3
β-1,3-(Glc)5
Na-acetate
MES
Na-acetate
MES
MES
PIPES
MES
PIPES
PIPES
MOPS
MOPS
HEPES
HEPES
TRIS
HEPES
TRIS
TRIS
pH 5.0
pH 5.5
pH 6.0
pH 6.5
pH 7.0
pH 7.5
pH 8.0
pH 8.5
pH 9.0
C
Quantification of thin layer chromatography
relative band intensity (%)
100
75
50
25
0
UT
30
40
50
60
70
80
temperature (°C)
sodium acetate
MES
sodium acetate
MES
MES
PIPES
MES
PIPES
PIPES
MOPS
MOPS
HEPES
HEPES
TRIS
HEPES
TRIS
TRIS
relative band intensity (%)
100
75
50
25
0
5
5.5
6
6.5
7
7.5
8
8.5
9
pH
other
glucose
laminaribiose
laminaritriose
> laminaripentaose

**Supplementary Figure 9. Temperature- and buffer-dependency of GBP1 activity on laminarin.** (A) Laminarin (4 mg·mL$^{-1}$) was digested with GBP1 (70 nM) for 10 min at different temperatures. Sample without enzymes (UT) was mock-digested at 80°C. (B) Laminarin (4 mg·mL$^{-1}$) was digested with purified GBP1 (70 nM) in different buffers (10 mM, pH 5-9). Digestions were performed at 60 °C for 10 min. Products from GBP1-catalyzed laminarin digestion assays in (A) and (B) were separated by thin layer chromatography. (C) Quantification of thin layer chromatography band intensities using the ImageJ software. UT, untreated sample.

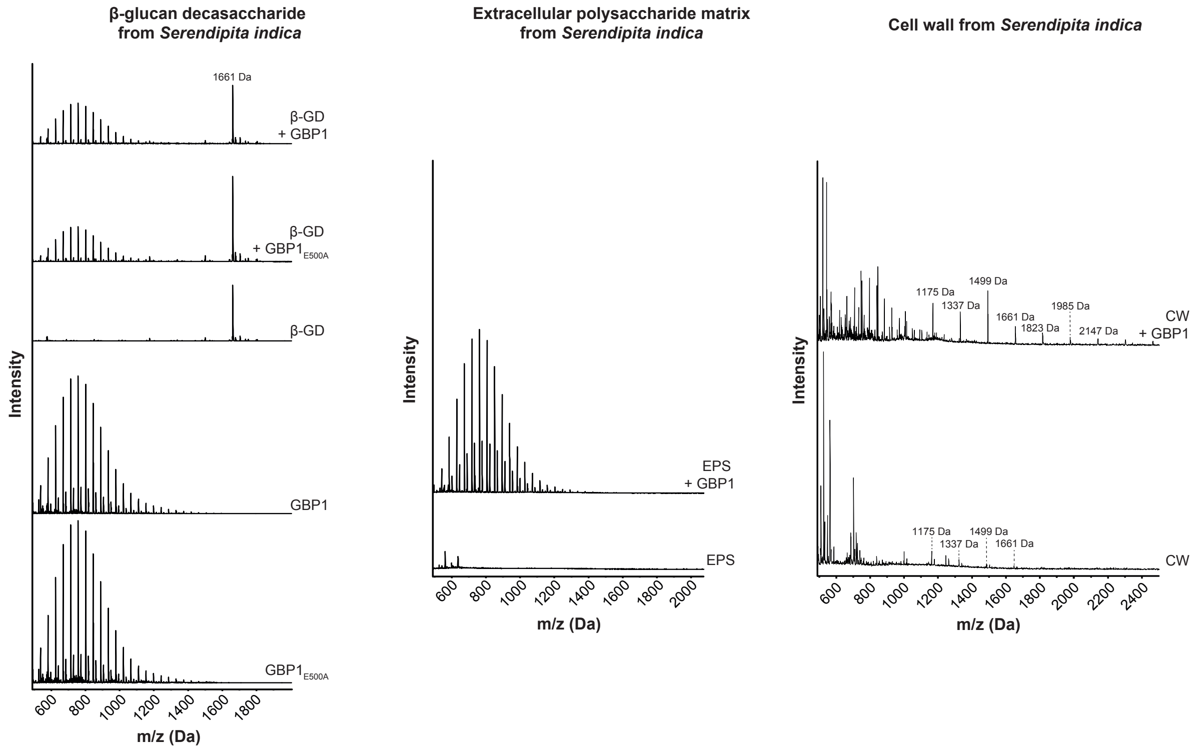


**Supplementary Figure 10.** Analysis of GBP1-digested β-glucan decasaccharide, extracellular polysaccharides, or cell wall fractions from *Serendipita indica* by MALDI-TOF mass spectrometry. While GBP1 does not act on the EPS matrix and the derived β-GD from *S. indica*, it releases minor oligosaccharide fractions [(m/z 1175 ($Hexose_7$) – m/z 1247 ($Hexose_{13}$)] from the CW. Oligosaccharide peaks of interest were labeled with *m/z* $(M+Na)^+$ masses. The digestion assays using the β-GD were performed two times with similar results and the digestion assays using the EPS and CW were performed once. β-GD, β-glucan decasaccharide; CW, cell wall; EPS, extracellular polysaccharides.

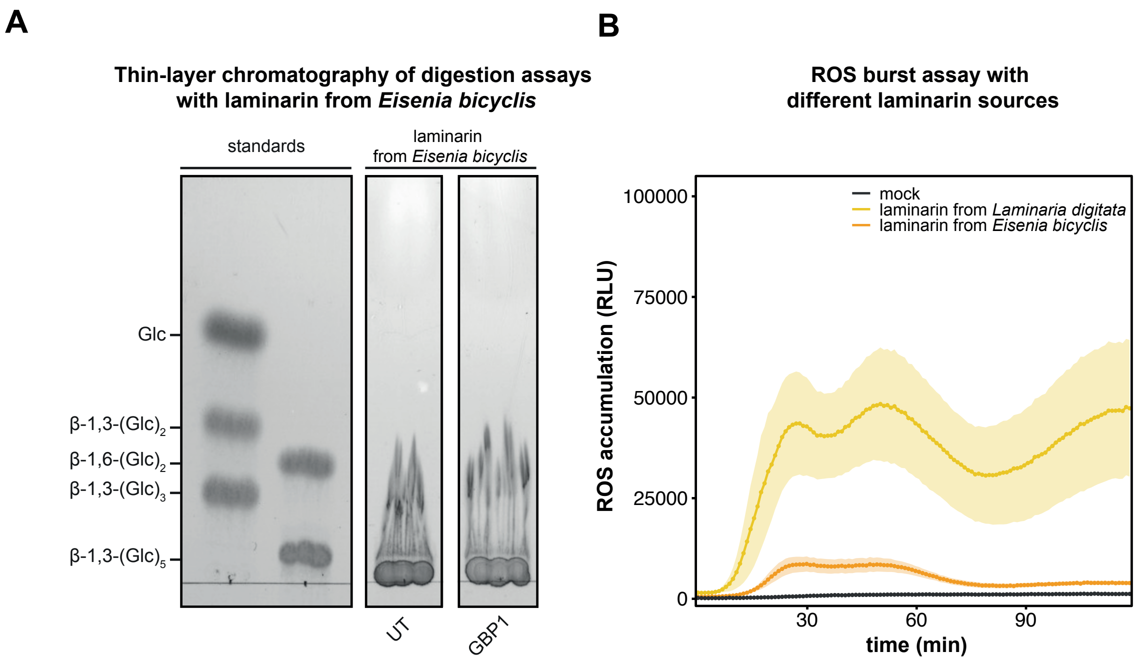


**Supplementary Figure 11. Highly branched laminarin from *Eisenia bicyclis* is inert to GBP1 hydrolysis and triggers only low production of ROS in barley roots.** (A) Laminarin from *E. bicyclis* (2.4 mM) was digested with GBP1 (72 nM) for 1 h at 60°C. The digestion products were analyzed by thin layer chromatography. (B) Apoplastic ROS accumulation after treatment of barley root pieces with laminarin from either *Laminaria digitata* (low frequency of β–1,6 linked branches) or *E. bicyclis* (high frequency of β–1,6 linked branches) was monitored by ROS burst assay. Treatment with Milli-Q water (mock) was used as control. Values represent mean ± SEM from eight wells. ROS, reactive oxygen species; RLU, relative luminescence units; UT, untreated.

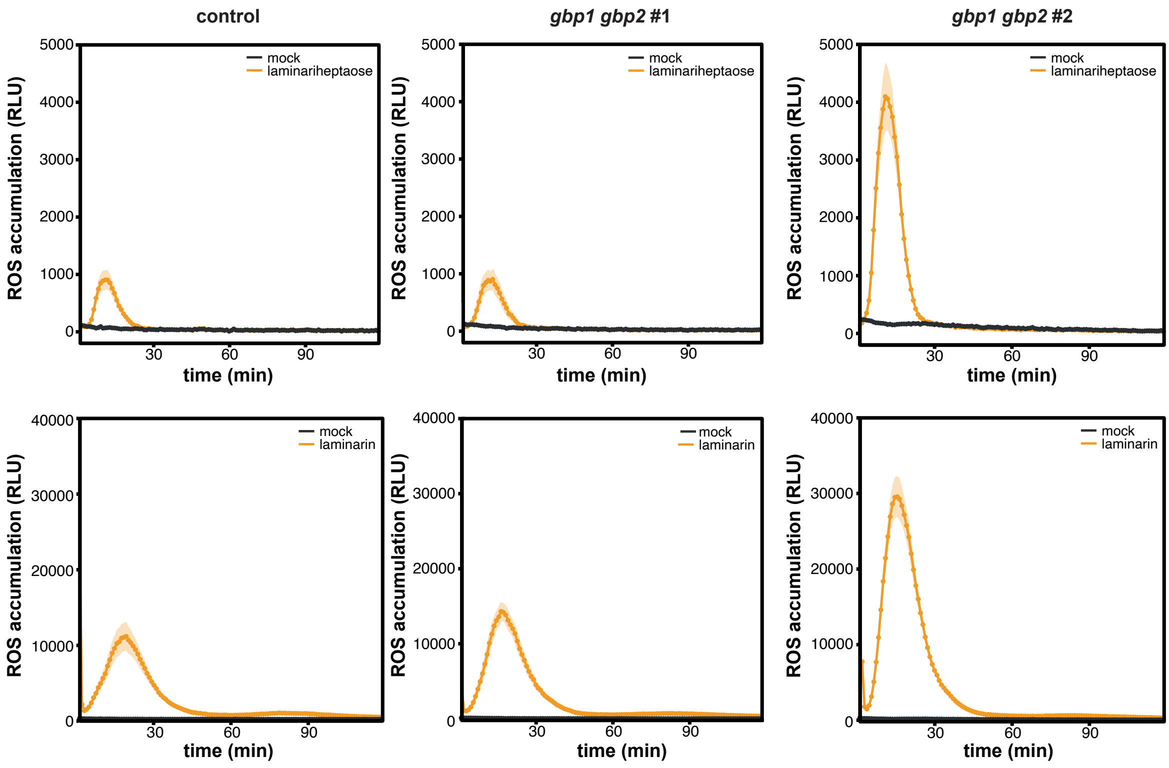


**Supplementary Figure 12. ROS burst assays in the barley control and *gbp1 gbp2* mutant lines.** Apoplastic ROS accumulation after treatment of barley roots with laminariheptaose (250 µM) and laminarin (4 mg·mL$^{-1}$) was quantified in control lines and *gbp1 gbp2* mutant lines. Treatment with Milli-Q water (mock) was used as control. Values represent mean ± SEM from 16 wells. The experiment was performed twice with similar results. ROS, reactive oxygen species; RLU, relative luminescence units.

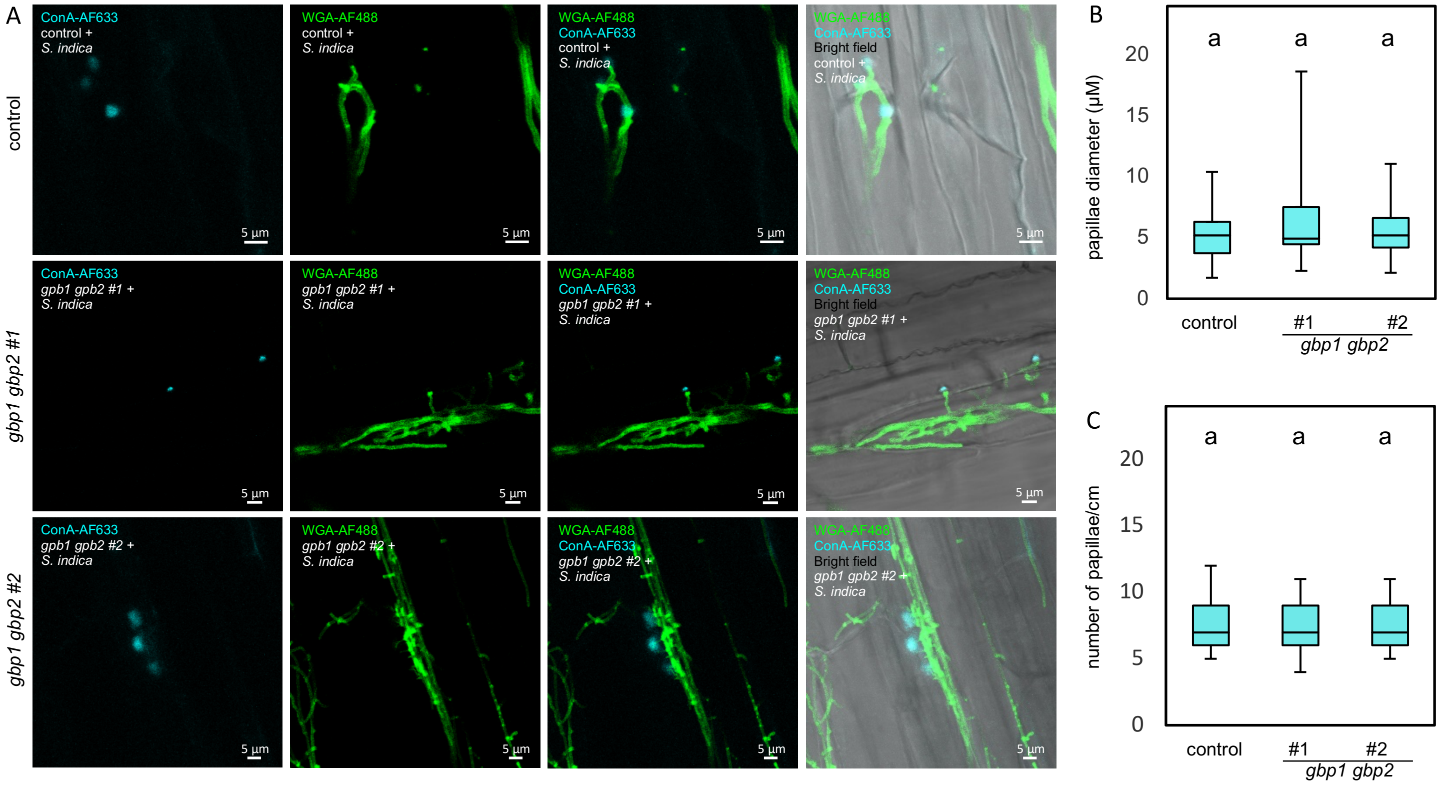


**Supplementary Figure 13. The number and size of papillae upon *Serendipita indica* colonization does not differ between the control and *gbp1 gbp2* mutant lines.** (A) *S. indica*-colonized roots of the barley control line and *gbp1 gbp2* mutant lines were stained with concanavalin A (ConA-AF633, cyan) and wheat germ agglutinin (WGA-AF488, green) for visualization of papillae and fungal structures, respectively. Images were acquired using a confocal microscope. (B) The diameter of papillae found in thecontrol and *gbp1 gbp2* mutant lines colonized by *S. indica*. (C) The number of papillae were quantified in thecontrol line and *gbp1 gbp2* mutant lines colonized by *S. indica*. ConA, concanavalin A; CW(A), cell wall (appositions); WGA, wheat germ agglutinin. Different letters represent statistically significant differences based on one-way ANOVA and Tukey's post hoc test with significance threshold: $P$ ≤0.05.

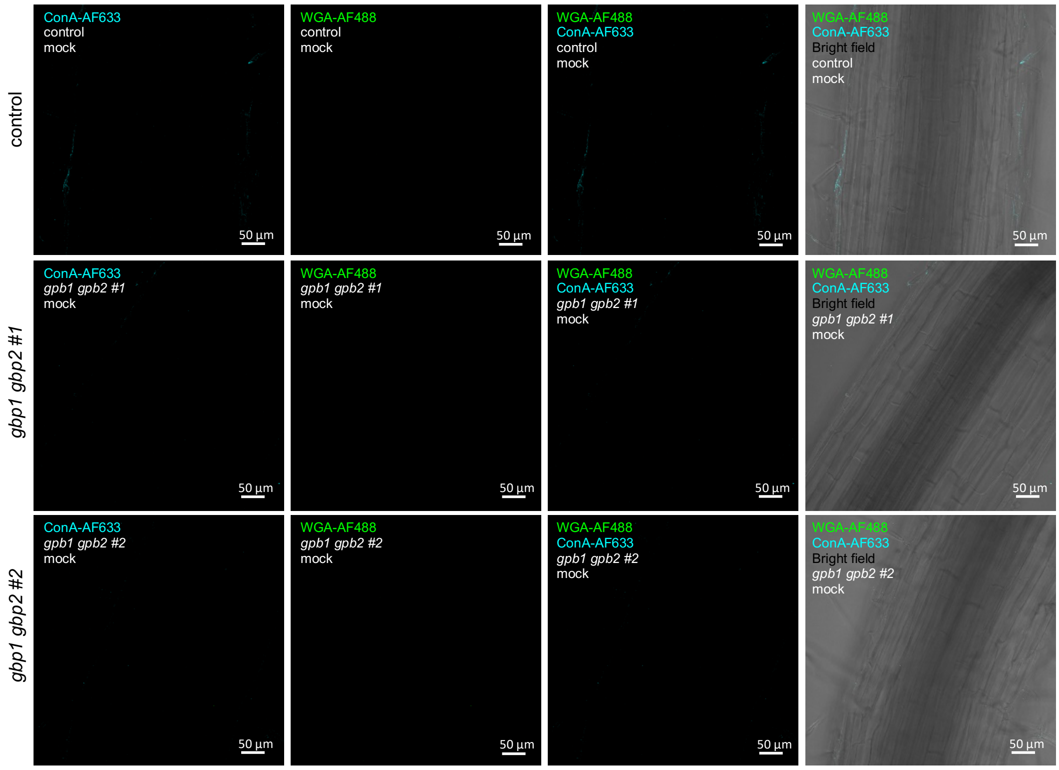


**Supplementary Figure 14. The barley control and *gbp1 gbp2* mutant lines do not form spontaneous papillae or CW responses in absence of fungal colonization.** Uninoculated roots of the control and *gbp1 gbp2* mutant lines were stained with concanavalin A (ConA-AF633, cyan) and wheat germ agglutinin (WGA-AF488, green) for visualization of CWAs and fungal structures, respectively. Images were acquired using a confocal microscope. ConA, concanavalin A; CW(A), cell wall (appositions); WGA, wheat germ agglutinin.
